# Structure and mechanism of oxalate transporter OxlT in an oxalate-degrading bacterium in the gut microbiota

**DOI:** 10.1101/2021.11.15.468502

**Authors:** Titouan Jaunet-Lahary, Tatsuro Shimamura, Masahiro Hayashi, Norimichi Nomura, Kouta Hirasawa, Tetsuya Shimizu, Masao Yamashita, Keiichi Kojima, Yuki Sudo, Takashi Tamura, Hiroko Iwanari, Takao Hamakubo, So Iwata, Kei-ichi Okazaki, Teruhisa Hirai, Atsuko Yamashita

**Author notes:** Corresponding authors: Tatsuro Shimamura, Graduate School of Medicine, Kyoto University, Kyoto, 606-8501, Japan.; Kei-ichi Okazaki, Research Center for Computational Science, Institute for Molecular Science, National Institutes of Natural Sciences, Okazaki, 444-8585, Japan.; Teruhisa Hirai, Current address: Japan Science and Technology Agency, Tokyo, 102-8666, Japan; Atsuko Yamashita, Graduate School of Medicine, Dentistry and Pharmaceutical Sciences, Okayama University, 1-1-1, Tsushima-naka, Kita-ku, Okayama 600-8530, Japan.

## Abstract

*Oxalobacter formigenes* is an oxalate-degrading bacterium in the gut microbiota that absorbs food-derived oxalate to use this as a carbon and energy source and thereby helps reduce the risk of kidney stone formation of the host animals ^1–4^. The bacterial oxalate transporter OxlT uptakes oxalate from the gut to bacterial cells and excrete formate as a degradation product, with a strict discrimination from other carboxylates that serve as nutrients ^5–7^. Nevertheless, the underlying mechanism remains unclear. Here, we present crystal structures of oxalate-bound and ligand-free OxlT in two different conformations, occluded and outward-facing states. The oxalate binding site contains two basic residues that form salt bridges with a dicarboxylate substrate while preventing the conformational switch to the occluded state without an acidic substrate, a ‘disallowed’ state for an antiporter ^8, 9^. The occluded ligand-binding pocket can accommodate oxalate but not larger dicarboxylates, such as metabolic intermediates. The permeation pathways from the binding pocket are completely blocked by extensive interdomain hydrophobic and ionic interactions. Nevertheless, a molecular dynamics simulation showed that a flip of a single side chain neighbouring the substrate is sufficient to trigger the gate opening. The OxlT structure indicates the underlying metabolic interactions enabling favourable symbiosis at a molecular level.

## Introduction

Oxalate is the smallest dicarboxylate (C_2_O_4_^2–^) ingested through our daily diet from oxalate-containing foods ^10^, such as vegetables, beans and nuts ^11^. Oxalate is also a final metabolic product in our body and is partly secreted to the intestine via the systemic circulation ^10^. Then it is absorbed from the intestinal tract and excreted through the kidney ^2^. However, excess oxalate forms an insoluble salt with blood calcium and causes kidney stone disease (Fig. 1A). *Oxalobacter formigenes* is an oxalate-degrading bacteria in the gut ^12^ that can degrade intestinal oxalate and thus significantly contribute to oxalate homeostasis in the host. Indeed, patients with cystic fibrosis ^13^ or inflammatory bowel disease ^14^ or those who have undergone jejunoileal bypass surgery ^15^ are known to have low rates of colonisation of *O. formigenes* and an increased risk of hyperoxaluria and kidney stone formation.

**Fig. 1.**
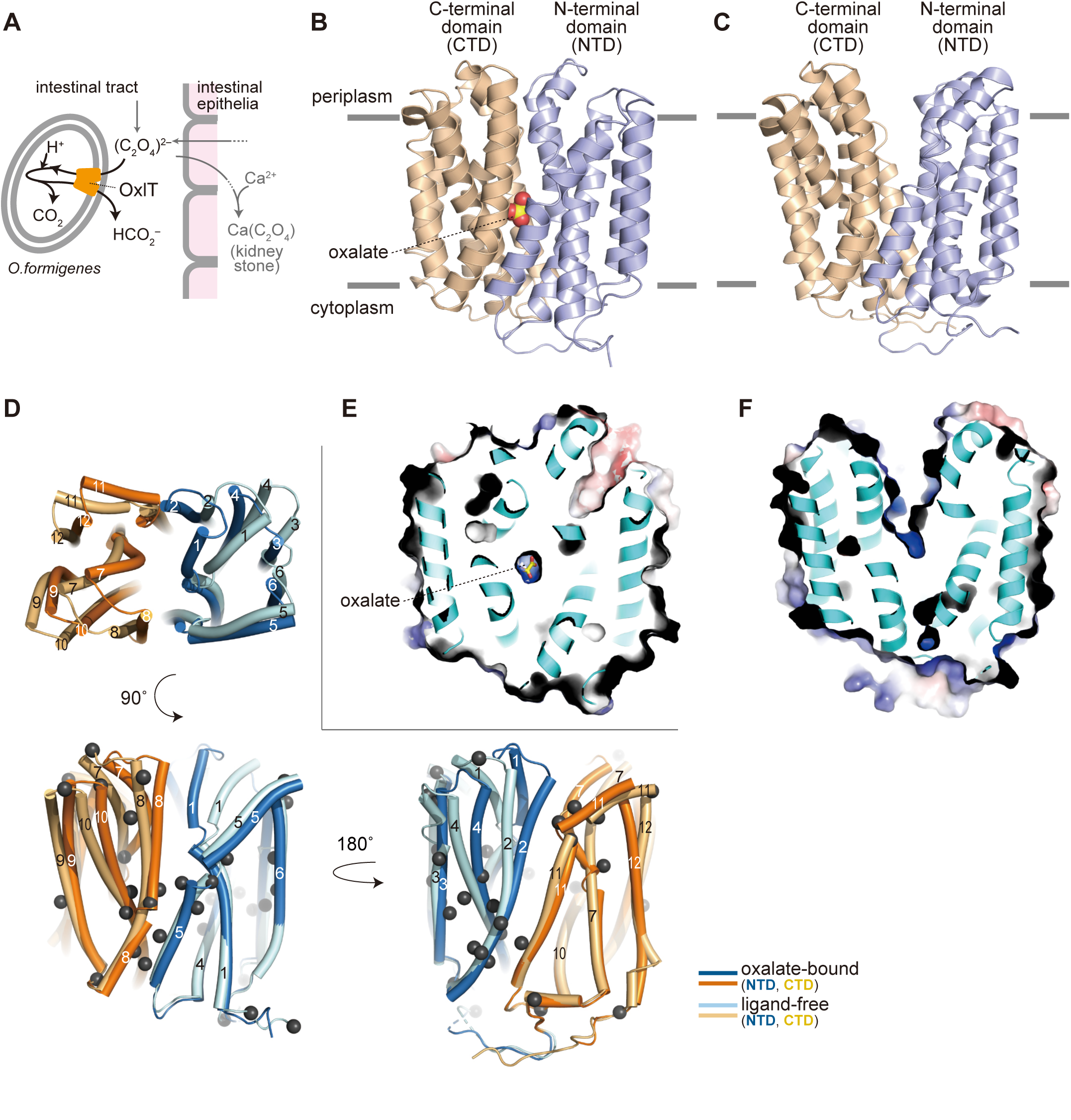
Structure of OxlT. (A) Schematic drawing of OxlT function in the oxalate-degrading bacterium, *O. formigenes*, in the gut. (B, C) Crystal structures of the oxalate-bound (B) and ligand-free (C) OxlT; (D) Superposition of oxalate-bound and ligand-free OxlT. A view from the periplasm (top) and two views in the transmembrane plane (bottom) are shown. Dark grey spheres indicate Cα atoms of glycine residues. (E, F) Surface electrostatic potential map of oxalate-bound (E) and ligand-free (F) OxlT. Electrostatic potentials at ± 5 kTe^-1^ were mapped on the surfaces.

Oxalate transporter (OxlT), an oxalate:formate antiporter (OFA) in *O. formigenes*, is a key molecule for oxalate metabolism in this bacterium. OxlT catalyses antiport of carboxylates across the cell membrane according to their electrochemical gradients with a substrate specificity optimised to the C2 dicarboxylate, oxalate. Indeed, the transporter shows a high turnover rate (>1000/s) for oxalate self-exchange ^5, 7^. Under physiological conditions in the oxalate autotroph *O. formigenes*, the carboxylate-exchange function of OxlT enables uptake of oxalate from the host intestine as a sole carbon source for the bacterium and a release of formate (HCO_2_^−^), the final degradation product of oxalate that is toxic if accumulated in the cell ^5–7^ (Fig. 1A). OxIT catalytic turnover of the oxalate:formate exchange is accompanied by the metabolic degradation of oxalate to formate via a decarboxylase that consumes a proton in the cytosol, consequently producing a proton electrochemical gradient across the bacterial cell membrane ^5^. Therefore, OxlT serves as a ‘virtual proton pump’ that creates a proton motive force for bacterial ATP synthesis ^5^. Thus, the functional characteristics of OxlT as an antiporter between oxalate and formate, rather than a uniporter of each chemical, is essential to couple carbon metabolism and energy formation. Notably, OxlT does not accept oxaloacetate or succinate, which are Krebs cycle dicarboxylate intermediates, as substrates ^6^. These dicarboxylates with four carbon atoms (C4 dicarboxylates) are important metabolic intermediates at the bacterial cytosolic side while they are also absorbed as energy sources and biosynthetic precursors through an intestinal transporter at the host lumen side ^16^. Therefore, the ability of OxlT to discriminate between C2 and C4 dicarboxylates is critical for the favourable symbiosis between host animals and the gut bacterium.

OxlT belongs to the major facilitator superfamily (MFS), the large transporter family whose members transport a wide array of chemicals ^17^. MFS proteins share a common architecture of twelve transmembrane (TM) helices that contain symmetrical N- and C-terminal halves of six gene-duplicated TM units, with a substrate-binding site in the centre of the molecule ^18, 19^. The substrate transport mechanism of the MFS family as well as other transporter families, is explained by the ‘alternating access model’ ^20, 21^, whereby transporter molecules open a cavity from the binding site to either side of the membrane alternately, and take outward-facing, occluded and inward-facing conformations via a ‘rocker switch’ motion of the N- and C-terminal domains, thereby allowing substrate transfer across the membrane ^22^. Although a wealth of structural information of each MFS member has been accumulated ^23–25^, current knowledge about the OFA family remains limited to an OxlT structure initially solved by electron crystallography at 6.5 Å ^8, 26^. Therefore, the specific oxalate recognition and antiport mechanism of OxlT is yet to be elucidated in a higher resolution structure. In this study, we report the X-ray crystallographic structures of OxlT in oxalate-bound and ligand-free forms solved at 3.0–3.3 Å to understand the structural basis of these key transporter functions that underly the symbiosis of this oxalate-degrading bacterium in the gut.

### OxlT structures in two different conformations

The wild-type OxlT is unstable under various conditions, such as in the presence of chloride ion ^7, 27^, which significantly narrows the available chemical space for crystallisation screening. Therefore, OxlT was stabilised by binding antibody fragments, resulting in crystallisation under two different conditions. We confirmed that the Fab fragment used for crystallisation binds to OxlT both in the presence and absence of oxalate, suggesting that Fab-mediated artificial trapping of OxlT in a certain conformation is unlikely. The crystal structure of oxalate-bound OxlT in complex with the Fab fragment was solved at 3.0 Å while that of ligand-free OxlT in complex with an Fv fragment was solved at 3.3 Å (Extended Data Fig. 1A, 1B, Extended Data Table 1).

The overall structure of OxlT consists of 12 TM helices (Fig. 1B, 1C), as observed in the previous EM structure ^26^ and later confirmed as a typical MFS architecture ^18, 19^. In the oxalate-bound state, the OxlT molecule adopts an occluded conformation with an oxalate molecule binding at the centre of the structure (Fig. 1, B and E). In contrast, the ligand-free OxlT takes a significantly different conformation from the oxalate-bound form (Fig. 1C, D, F). The OxlT molecule displayed a large V-shaped cavity between the N- (TM1–6) and the C-terminal (TM7–12) domains, which was connected from the central oxalate binding site to the periplasm, a clear signature of an outward-facing conformation.

In a comparison of the occluded and outward-facing structures, the Cα root-mean-square-deviation (RMSD) for all residues was 2.6–2.7 Å (Fig. 1D). Even the sole N- or C-terminal domains of the two showed significant structural differences (Cα RMSD of 1.5∼1.6 Å). Therefore, the structural change between the occluded and outward-facing states with a ‘rocker switch’ motion is not achieved by the tilt of the rigid structural units but is concomitant with their bending. Indeed, conspicuous bends at the periplasmic portion were observed on TM1, 2, 4, 7, 8 and 11 in the outward-facing structure, with tilting of the other surrounding TM helices (Fig. 1D). In contrast, there was no significant change of the cytoplasmic portion between the two conformations. In other MFS proteins, such as GLUT5 and NarK, bending at the glycine residues in the TM helices has been observed between the different conformational states ^28, 29^. OxlT has 52 glycine residues, which is one eighth (12.4%) of the amino acid content (Fig. 1D, Extended Data Fig. 1C, Extended Data Fig. 2). Notably, this glycine frequency is significantly higher than that in other MFS proteins, such as LacY (8.6%), GLUT5 (7.6%) or NarK (10.4%), and in TM helices in other membrane proteins (∼8.7%) ^30^. Therefore, the accumulation of bending of the TM helices at the glycine residues is likely more prominent in achieving the conformational switch between the states in OxlT. Glycine residues were also found at the interface between the N- and C-terminal domains as in TM5 and TM8 or TM2 and TM11 (Extended Data Fig. 1C) and achieved tight helical packing as previously reported ^30, 31^. The high glycine occurrence observed in OxlT may be required to occlude the oxalate, which is small for a transported substrate, in the centre of the molecule.

**Fig. 2.**
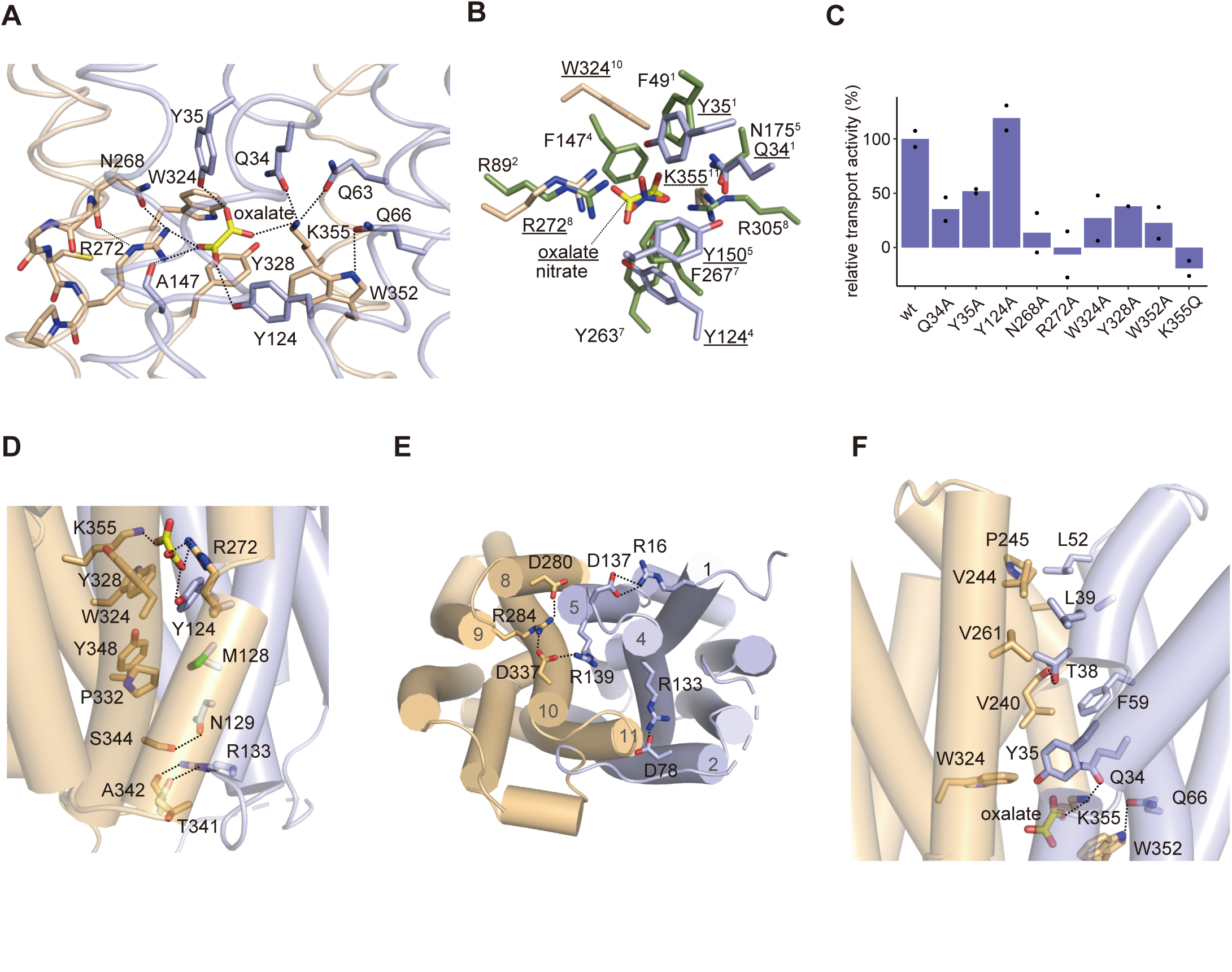
Oxalate-bound occluded OxlT structure. (A) Close up of the binding site in oxalate-bound OxlT. The domain colour coding is as in Fig. 1B. Dashed lines indicate potential hydrogen or ionic bonds. (B) Superposition of the substrate-binding site structures of OxlT and NarK (PDB ID: 4U4W) based on the topological similarity of the amino acid residues interacting with the substrates. OxlT is shown in the same colour coding as panel A with the underlined labels while NarK is shown in green with normal labels. The superscript for a residue label is the TM helix numbering where the residue locates. (C) Transport activities of the mutant OxlT relative to that of wild-type OxlT. The results of the R272A and K355Q mutants ^37^ are reposted for comparison. (D) Interdomain interactions closing the cavity to cytoplasm. (E) Ionic interaction network at the cytoplasmic side of OxlT. (F) Interdomain interactions closing the cavity to periplasm.

### Oxalate-bound occluded structure

In the crystal structure, the oxalate molecule binding to OxlT refined as a twisted configuration (Fig. 2A, Extended Data Fig. 3A). The bond between the two carboxyl groups in an oxalate dianion is known to be a single and unconjugated, allowing a free rotation of the carboxyl groups about the C-C bond ^32^. Since the resolution of the oxalate-bound OxlT crystal is insufficient to accurately determine the dihedral angle of oxalate, we performed QM and QM/MM calculations of the oxalate binding in the occluded OxlT structure to examine the energetically minimised conformation. The resulting O-C-C-O dihedral angles in the oxalate were within 50–68° (Extended Data Fig. 3B, 3C, Extended Data Table 2). These values are close to those observed in the original crystal structure (60.1°), verifying that the oxalate is not planar but twisted in the crystal structure.

**Fig. 3.**
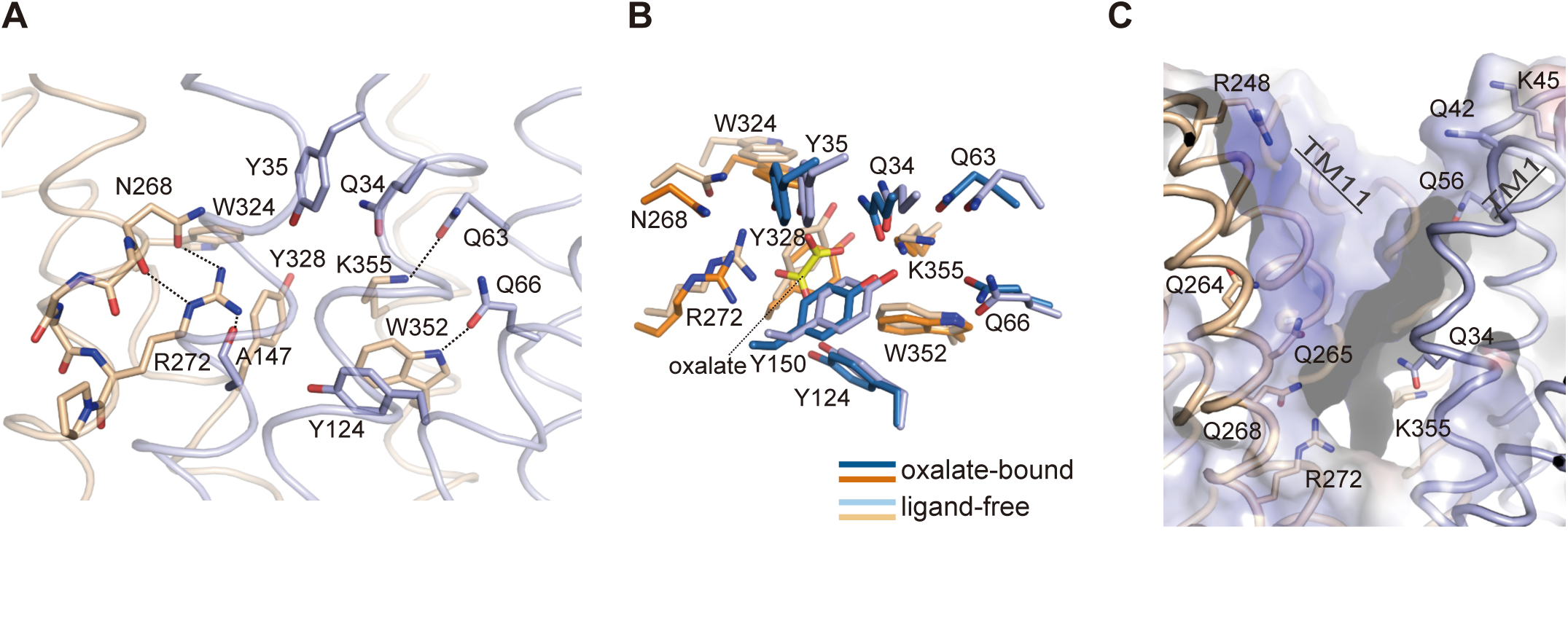
Ligand-free outward-facing OxlT structure. (A) Close up of the binding site in ligand-free OxlT viewed from the same orientation in Fig. 2A. The domain colour coding is as in 1C. The dashed lines indicate potential hydrogen bonds. (B) Superposition of the substrate-binding site structures of OxlT in oxalate-bound and ligand-free forms. (C) Close up of the cavity open to the periplasm. Models of polar residues exposed to the cavity and the surface coloured with the electrostatic potential map as in Fig. 1F are also shown. In panels A-C, the molecule defined as chain A is shown as a representative.

At the binding site in OxlT, oxalate binds to the transporter with one carboxyl group forming a bidentate salt bridge with Arg272 in TM8 while the other forms an ionic interaction with Lys355 in TM11 (Fig. 2A). In addition to the salt-bridging with the oxalate, the *ε*-amino group in Lys355 forms an interdomain hydrogen bond network with the carbamoyl groups in Gln34 (TM1) and Gln63 (TM2) in the N-terminal domain. Similarly, the guanidino group in Arg272 forms an interdomain hydrogen bond with the main chain carbonyl group in Ala147 (TM5) and further interacts with the carbamoyl and main chain carbonyl group of Asn268 upstream of TM8. The region around Arg272 is the bending point in TM8 due to the sequence of N^268^GGCR^272^P, and therefore the hydrogen bonds between Arg272 and Asn268 likely maintain the conformation and orientation of TM8 in the oxalate-bound structure. These inter- and intra-domain hydrogen bonding networks involving Arg272 and Lys355 likely play pivotal roles in organising the structure of the binding pocket and stabilising the occluded conformation, despite the location of these two basic residues within the C-terminal domain. These two basic residues are critical for oxalate transport, and even R272K or K355R mutations decrease the transport activity ^33–35^. These results confirm observations that not only the charges but also the chemical structures of the side chains of the two residues are important for the structural organisation of the binding site.

In addition to the two basic residues, numerous aromatic residues are found to contribute to oxalate binding. The hydroxyl groups of Tyr35 and Tyr124 form hydrogen bonds with either of the carboxyl groups in oxalate (Fig. 2A). Furthermore, the aromatic side chain groups in Tyr150, Trp324, Tyr328 and Trp352 form face-to-face or edge-to-face π-π interactions with the carboxyl groups in oxalate, indicating the significance of the π-electron systems in oxalate for molecular recognition by OxlT. These aromatic residues distributed in both the N- and C-terminal halves, and thus their interactions with oxalate, are also significant in stabilising the closure of the interdomain cavities to achieve the occluded conformation. In addition, Trp352 forms an interdomain hydrogen bond with Gln66 (TM2). Notably, a similar combination of ionic and π-π interactions was also observed with the recognition of nitrate, which also presents a π-electron system, by NarK in the nitrate/nitrite porter (NNP) family ^28, 31^, although the NNP family is distant from the OFA family ^17^ and the positions of the involving residues do not correspond to each other (Fig. 2B).

The significance of the above-mentioned or their neighbouring residues for oxalate transport was verified by a functional assay using *E. coli* recombinant expressing wild-type and mutant OxlT. In the assay, the extent of oxalate-formate exchange by OxlT, which is negatively electrogenic, was assessed by coupling with light-driven inward proton transfer by a microbial rhodopsin, xenorhodopsin ^36^, co-expressed in *E. coli* ^37^. In addition to R272A and K355Q, which are the reported non-functional mutants ^33, 34^ and had been verified their loss of activity in our previous study ^37^, mutation of OxlT with Q34A, Y35A, N268A, W324A, Y328A and W352A also reduced activity, indicating the significance of these residues for transport function (Fig. 2C). These residues are conserved among the OFA family proteins (Extended Data Fig. 2).

The interactions between the substrate and the residues in the occluded OxlT crystal structure are optimised to the C2 dicarboxylate oxalate. Since the oxalate molecule tightly fits to the binding pocket, supplanting oxalate with a larger dicarboxylate, such as Krebs cycle intermediates, causes steric clashes with the residues in OxlT, likely destabilising the occluded conformation (Extended Data Fig. 4A). A flexible docking study resulted in a position that could accommodate a C3 dicarboxylate, malonate, in the binding site of the occluded OxlT, although this had fewer interactions compared with the case for oxalate, due to the rearrangement of amino acid residues in the binding pocket (Extended Data Fig. 4B). This is consistent with the reduced affinity and transport activity to malonate ^6^. No pose for the binding of C4 dicarboxylates to the occluded OxlT was observed even by flexible docking, consistent with a prior report indicating that there was no significant binding of these molecules to OxlT ^6^.

**Fig. 4.**
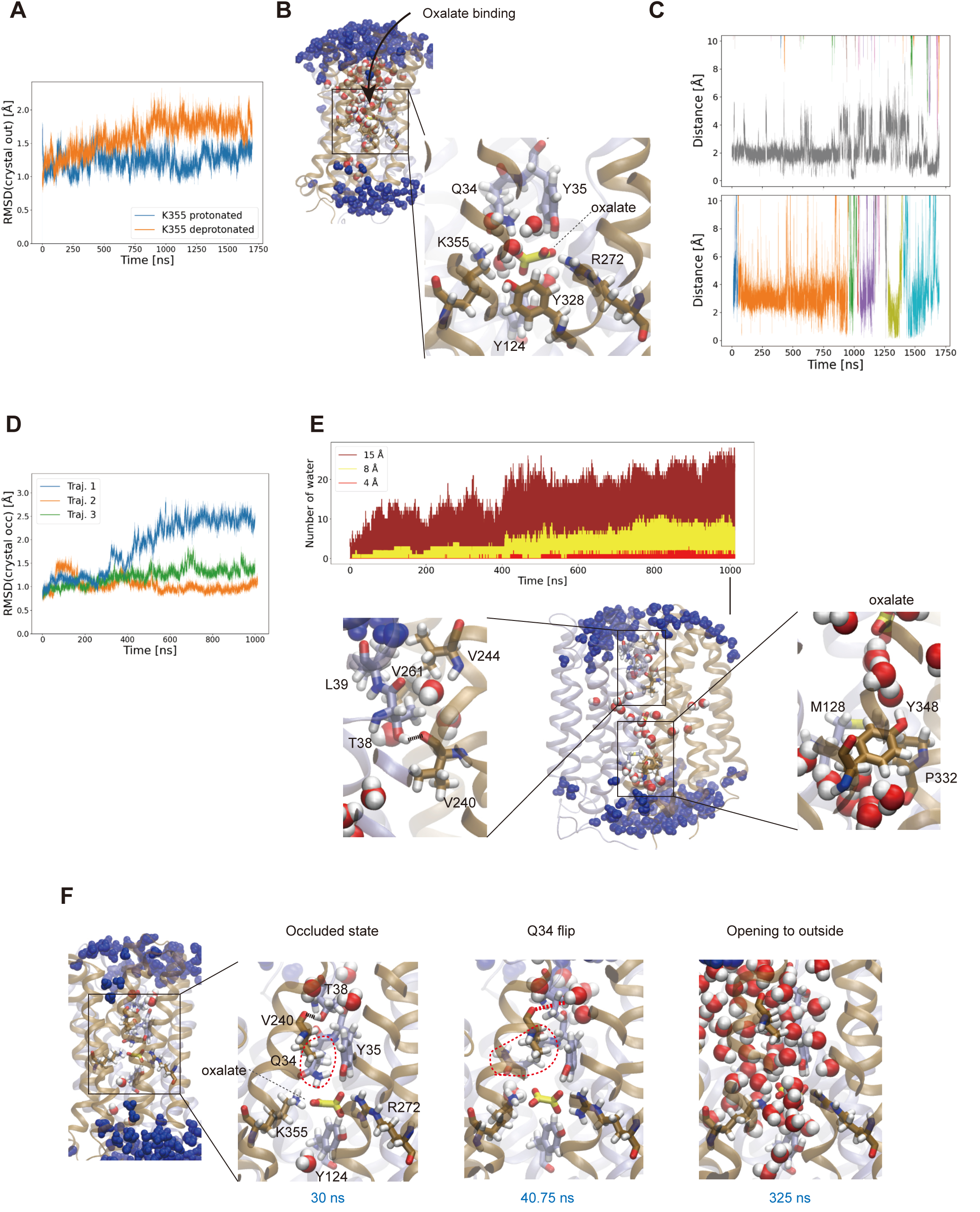
Substrate-binding and conformational dynamics of OxlT. (A–C) MD simulations started from the ligand-free outward-facing OxlT crystal structure. (A) RMSDs from the initial outward-facing crystal structure are shown for two trajectories with different protonation states of Lys355 in different colours. (B) Snapshot of the spontaneously bound oxalate to OxlT with protonated Lys355. In the zoom-out snapshot, water molecules within 15 A of the oxalate ion are in the CPK colour while those between 15 and 25 A are in blue. In the close up snapshot, water molecules within 4 A distance from the oxalate ion are shown. (C) Time series of the distance between the oxalate ions and the binding site residues with protonated and deprotonated Lys355 are in the top and bottom panels, respectively. Different colours represent different oxalate ions. (To be continued.) (D–F) MD simulations started from the oxalate-bound occluded OxlT crystal structure. (D) RMSDs from the initial occluded crystal structure are shown for three independent trajectories in different colours. (E) Hydrophobic gates of OxlT. Top panel: the numbers of water molecules within 15, 8 and 4 Å from the bound oxalate ion are plotted in brown, yellow and red, respectively. Bottom panel: a snapshot at 1000 ns is shown in the zoom- out and close up views. Water molecules within 15 Å are in the CPK colour while those between 15 and 25 Å are in blue. (F) The observed transition from the occluded to the outward-facing conformation triggered by the Gln34 flip. The oxalate ion and binding site residues are represented as sticks. Gln34 is highlighted with the red circle. Water molecules are shown in the van der Waals representation. CPK-coloured water molecules are within 15 Å from the oxalate ion while the blue ones between 15 and 25 Å from the oxalate ion. The broken lines between the Thr38 side chain and the Val240 main chain in black and red depict the distances those within or out of H-bonding, respectively.

From the oxalate binding site to the cytoplasm or periplasm, extensive intramolecular interactions were observed between TM helices in the N- and the C-terminal domains, such as TM2 and TM11, TM5 and TM8, the periplasmic halves of TM1 and TM7, and the cytoplasmic halves of TM4 and TM10 (Fig. 2, D–F). These interactions stabilise the closure of the interdomain cavities in the occluded structure.

In the cytoplasmic side below the oxalate binding site, hydrophobic interactions involving Met128 (TM4), Pro332 (TM10) and Tyr348 (TM11) were observed, followed by polar interactions between Asn129 (TM4) and Ser344 (TM11), Arg133 (TM4) and the main chain carbonyl groups of Thr341 and Ala342 (TM11) (Fig. 2D). These interactions are further supported with by charge-dipole interactions at the cytoplasmic end, formed between Asp78 (TM2) or Asp280 (TM8) and the N-terminal ends of TM11 or TM5, respectively (Fig. 2E). The two aspartate residues located in the ‘A-like’ motifs in the TM2-3 (‘G^74^YFVD^78^KFGP^82^R^83^IP’ sequence, A^L^^2–3^) or TM8-9 (‘G^276^FVSD^280^KIGR^284^YK’, sequence, A^L^^8–9^) regions (Extended Data Fig. 2). Motif A is one of the commonly conserved motifs in MFS proteins, and the D(+5) is known to participate in an interdomain charge-helix dipole interactions ^9^. Notably, these aspartate residues further compose extensive ionic interaction networks in the cytoplasmic side (Fig. 2E). Specifically, Asp78 and Arg133 in TM4, and the downstream residue Asp137 and Arg16 in TM1, form salt bridges. Further downstream, Arg139 at the N-terminal end of TM5 forms a charge relay network with Asp337, Arg284 and Asp280.

In the periplasmic side above the oxalate binding site, a hydrogen bond between Thr38 (side chain) in TM1 and Val240 (backbone) in TM7 (2.72 Å) closes the pore tunnel in the occluded conformation (Fig. 2F). Above the hydrogen bond, Leu39 (TM1), Leu52 (TM2), Val244 and Pro245 (TM7) and Val261 (TM8) form hydrophobic interactions.

### Ligand-free outward-facing structure

In contrast to the occluded substrate-binding site in the oxalate-bound OxlT, a large cavity from the binding site to the periplasmic space is open in the ligand-free OxlT (Fig. 1F). At the empty binding site, the Lys355 side chain flips out from Arg272 due to charge repulsion and shifts the positions from those found in the oxalate-bound form (Fig. 3A). In the ligand-free form, most of the interdomain hydrogen bonds observed in the oxalate-bound state are retained. However, that between Lys355 in the C-terminal domain and Gln34 in the N-terminal domain is likely disrupted in the ligand-free state, judging by the distance between the side chains (>∼4 Å). Positional shifts of the surrounding aromatic residues, such as Tyr35, Tyr150, Trp324 and Tyr328, were also observed (Fig. 3B). These changes at the substrate-binding site due to the absence of oxalate likely underlie the structural rearrangement of the overall architecture and result in the conformational change between the occluded and outward-facing state. Notably, the cavity opening to the periplasm displayed an extensive positively charged surface (Fig. 1F, 3C). This basic property is mainly derived from Arg272 and Lys355 in the binding site. In addition, the side-chain amino groups in Lys45 and Arg248 and the amide groups in Gln34, Asn42, Gln56, Asn264, Asn265 and Asn268, that line this cavity, are now exposed to the solvent. These groups and the positive dipole moments of the bent helices of TM1, TM5 and TM11 also contribute to the basic property of the entire cavity (Fig. 3C). The charge repulsion caused by Arg272 and Lys355 at the empty ligand-binding site as well as the extensive basic surface of the cavity likely prevents closure of the pocket to the occluded form in the absence of oxalate, thus stabilising an open state. The stability of an open state conformation in the absence of a substrate, which prevents transition to the occluded state, underlies the OxlT function as an antiporter, in which the conformational switch in the absence of a substrate during the catalytic process is disallowed ^8, 9^. A similar situation was observed on a nitrate/nitrite antiporter NarK ^31^, where the positively charged surface of the open cavity stabilised the inward-facing conformation ^28^.

On the other hand, the cytoplasmic part of the ligand-free OxlT structure shows no significant changes from that of the oxalate-bound structure (Fig. 1D).

### Substrate-binding, hydrophobic gates and conformational dynamics of OxlT

To address the structural dynamics of OxlT enabling the conformational switch necessary for the transport cycle, we performed molecular dynamics (MD) simulations ^38^ based on the oxalate-bound occluded and the ligand-free outward-facing OxlT crystal structures.

We first simulated oxalate binding to the ligand-free outward-facing conformation (Fig. 4, A–C). Spontaneous binding of oxalate ion to the binding site of OxlT was observed at Gln34, Tyr35, Arg272, Tyr328 and Lys355 (Fig. 4B). An extensive positively charged surface (Fig. 1F, 3C) contributes to a rapid spontaneous binding of the negatively charged oxalate ion. The stability of the bound conformation was dependent on the protonation state of Lys355 (see Methods for p*K*_a_ calculation), which could be affected by the luminal pH in the gut, varying ∼ 5 – 8 by regions ^39^. For the protonated Lys355, a single binding event was observed, and the bound oxalate ion remained in the binding site for the rest of the simulation, whereas several binding and unbinding events were observed for the neutral Lys355 (Fig. 4C). During the 1.7 µs simulations, the outward-facing conformation of OxlT was stable, as shown in the plot of the RMSD of the backbone atoms from the outward-facing crystal structure (Fig. 4A).

The results suggest that the spontaneous binding of the oxalate observed in the simulations is an early-stage binding mode that should be followed by the conformational rearrangement and desolvation of the binding site and the transition to the occluded conformation to adapt the fully bound conformation.

We next addressed the conformational dynamics of the occluded conformation in the oxalate-bound state. During the 1 µs simulation, two out of three independent trajectories remained in the occluded state (Fig. 4D). In the occluded conformation, most water molecules were blocked at certain positions in the periplasmic and cytoplasmic sides of the transporter during the simulations, although some entered OxlT (Fig. 4E). A water density analysis pinpointed structural layers blocking entry of water into the oxalate binding site during the simulation (Extended Data Fig. 5). One of these is a hydrophobic layer constituting of Thr38 and Leu39 in TM1, Val244 in TM7 and Val261 in TM8 at the periplasmic side (lower left panel of Fig. 4E). This layer, combined with the hydrogen bond between Thr38 and Val240 in TM7 (shown by a broken line in lower left panel of Fig. 4E), also blocked the exit of ligand to the extracellular side and thus served as the periplasmic gate. The other layer consists of Met128 in TM4, Pro332 in TM10 and Tyr348 in TM11 at the cytoplasmic side (lower right panel of Fig. 4E). These periplasmic and cytoplasmic hydrophobic gates, together with the TM1–TM7 hydrogen bond, have similarity with the previously reported NarK transporter ^40^, based on residues located at similar positions to those in OxlT in the aligned structure (Extended Data Fig. 6). This result suggests that the hydrophobic gates ^38^ are a conserved mechanism among the two transporters.

**Fig. 5.**
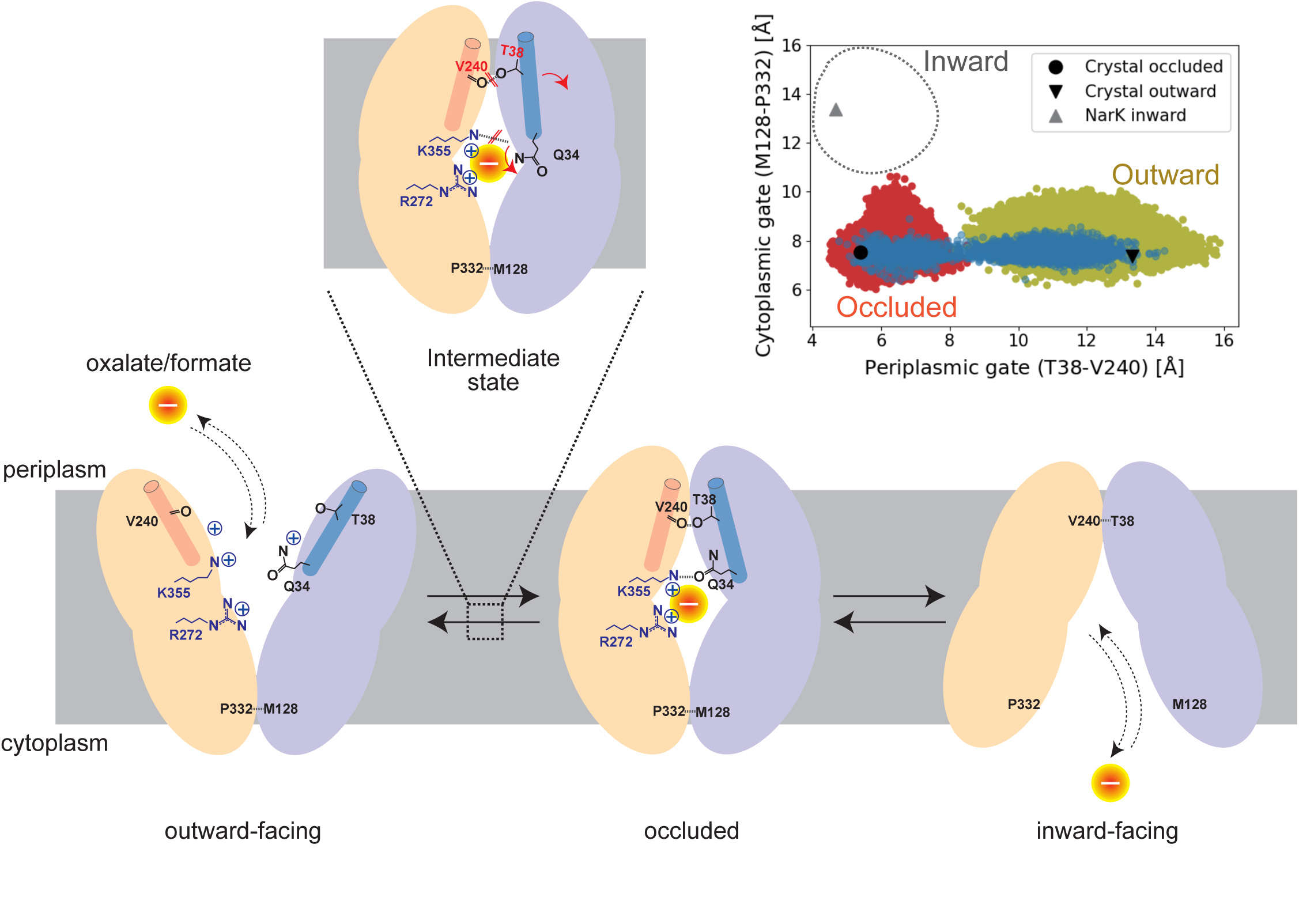
Schematic drawing of the transport process and conformational switching of OxlT. The conformational landscape of OxlT along the periplasmic and cytoplasmic gate distances is shown in the top right panel. The Cα distances of the gate residues in MD simulations of the occluded and outward- open states and Gln34-induced transition are shown in red, yellow and blue, respectively. The gate- residue distances in the current occluded and outward-facing crystal structures as well as the Nark inward-facing crystal structure (PDB ID: 4U4T) are also shown.

In contrast, in one trajectory from the occluded conformation, an opening of the periplasmic gate was observed (blue line in Fig. 4D). In the transition, the flip of Gln34 side chain from the binding site occurred first (Fig. 4F, Extended Data Fig. 7A). The Gln34 flip resulted in a disruption of the hydrogen bond with Lys355, as observed in the outward-facing crystal structure (Fig. 3A). Furthermore, since Gln34 is located one-turn upstream of Thr38 in TM1, the flip also caused a constant disruption of the hydrogen bond between Thr38 and Val240, which was bonded on and off by thermal fluctuation even before the flip (shown by a broken line in Fig. 4F; Extended Data Fig. 7B). After ∼280 ns following the Gln34 flip, OxlT started opening to the outside and many water molecules entered the transporter (Fig. 4F). We note that the Gln34 flip is a transient conformation and that the side chain returned to the original position after reaching the outward-facing state in the last part of the simulation, consistent with the observation on the outward-facing crystal structure (Extended Data Fig. 7A and 7C). Notably, the Gln34 flip was also observed in a trajectory starting from the occluded conformation with formate modelled in the binding site (Extended Data Fig. 8). In this trajectory, the hydrogen bond between Thr38 and Val240 was again completely broken after the Gln34 flip, followed by a transition from the occluded to the outward-facing conformation, in accordance with the physiological reaction of the formate release to periplasm in *O. formigenes*. In contrast, the Gln34 flip was not observed in any of the other trajectories unaccompanied with the conformational transition in both the oxalate- and formate-bound forms (Extended Data Fig. 7, 8). These results suggest that the Gln34 side chain, together with the hydrogen bond between Thr38 and Val240, works as a switch of the transition from the occluded to the outward-facing conformations. Indeed, Gln34 was identified as critical in the transport assay (Fig. 2C) and, together with Thr38, is strictly conserved among the OFA family (Extended Data Fig. 2).

The O-C-C-O dihedral angle of the oxalate ion in the occluded binding site became ∼90° after the Gln34 flip (Extended Data Fig. 9A), which is the value observed in solution ^41^. This contrasts with the other two trajectories without the Gln34 flip, where the oxalate dihedral angle remained around 40–50° (and its inverted position at 130–140°; Extended Data Fig. 9A), which is similar to those found in the crystal structure and the QM/MM calculations. Intriguingly, the values observed in the bound oxalate to the outward-facing OxlT during the simulation were broadly distributed with double peaks at ∼60° and ∼120° (Extended Data Fig. 9B), which differ from those in solution and rather closer to those in the occluded crystal structure. These results imply that the bound oxalate rearranges its conformation according to the environmental change derived by OxlT conformational changes and takes a favourable conformation to the next step in the transporter cycle.

No opening of the cytoplasmic gate was observed during the 1 µs simulation for any of the trajectories from the occluded conformation. This may be attributed to the extensive interdomain interactions observed at the cytoplasmic side, such as the motif A involving charge relay networks (Fig. 2E) known to stabilise the outward-facing conformation^9^. These results suggested that the transition from the occluded to the inward-open state has a slow kinetics among the entire transport process.

## Discussion

The two crystal structures of OxlT and the MD simulations based on them provided clues to understand the alternating access transport process of OxlT (Fig. 5). The following process is described according to the electrochemical gradient formed in *O. formigenes* within the gut. For the oxalate uptake process, OxlT exhibits an extensive positively charged surface in the cavity open to the periplasm, allowing a spontaneous binding of acidic oxalate to the binding site. The positively charged surface also avoids the conformational transition to the next transport step in the absence of the substrate that is an indispensable characteristic for an antiporter. Nevertheless, the oxalate binding neutralises the local positive charge and enables the conformational switch from the outward-facing conformation to the occluded conformation. The occluded state is an essential step for transport to serve as a discriminatory checkpoint between oxalate and necessary host metabolic intermediates, such as those in the Krebs cycle, using the size restriction imposed by the binding pocket. The occluded conformation may eventually allow opening of the cytoplasmic gate and release of oxalate to the cytoplasm.

Subsequently, a formate binding to the inward-facing OxlT may return the transporter to the occluded state. The conformational transition required for returning from the occluded to the initial outward-facing states in the antiport cycle can be achieved by a transient flip of a side chain of a substrate-neighbouring residue, Gln34, and disruption of the hydrogen bond between Thr38 and Val240. The conformational landscape plotted by the periplasmic gate (Thr38–Val240) and cytoplasmic gate (Met128–Pro332) distances ^40, 42^ sampled in the MD simulations shows that the order parameters separate the occluded and outward-facing conformations well (red and yellow plots, Fig. 5 inset). Nevertheless, the only trajectory that accompanies the Gln34 flip shows a full transition covering the endpoint occluded and outward-facing crystal structures (blue points in Fig. 5 inset).

These structural observations imply that OxlT utilises the MFS architecture and evolved in accordance with favourable symbiosis between the host animals and gut microbes. The structural and functional characteristics of OxlT also likely underlie those of the other OFA family members. Approximately 2000 OFA members are registered in the database ^43^, and all but OxlT are functionally uncharacterised. Therefore, knowledge concerning OxlT also contributes to understanding unknown ‘dark’ protein families. Clarifying the inward-facing conformations of OxlT (a dotted circle in Fig. 5 inset) is the next challenge understanding the structural biology of OxlT.

## Methods

### Preparation of OxlT

C-terminal nona-His-tagged OxlT was expressed in *E. coli* XL3 at 20 °C for 24 h with 1 mM isopropyl-β-D-thiogalactopyranoside (IPTG) ^44^. Bacterial cells were suspended in lysis buffer (50 mM Tris-HCl, 200 mM K acetate, 1 mM EDTA, 1 mM PMSF, 5 mM MgCl_2_, 20 µg/mL DNaseI and 0.23 mg/mL lysozyme) and then disrupted using EmulsiFlex C-5 (Avestin). Cell debris was removed by centrifugation (9,600 × *g* for 30 min), and cell membranes were then collected by centrifugation (185,000 × *g* for 1 h). The membrane fraction was solubilised with 40 mM dodeclymaotoside (DDM) in buffer A (20 mM HEPES-KOH, 200 mM potassium acetate, 10 mM potassium oxalate and 20% glycerol, pH 8.0) and applied to Ni-NTA Superflow resin (QIAGEN) or HisTrap FF crude (GE Healthcare) in an XK16 column (GE Healthcare). The column was washed with buffer A (1 mM DDM and 30–50 mM imidazole), and then protein was eluted with buffer A containing 1 mM DDM and 250 mM imidazole.

### Preparation of antibody fragments

All animal experiments conformed to the guidelines of the Guide for the Care and Use of Laboratory Animals of Japan and were approved by the Animal Experimentation Committee at the University of Tokyo. A proteoliposome antigen was prepared by reconstituting purified functional OxlT at high density into phospholipid vesicles consisting of a 10:1 mixture of egg phosphatidylcholine (PC) (Avanti Polar Lipids) and adjuvant lipid A (Sigma) to facilitate an immune response. BALB/c mice were immunised with the proteoliposome antigen using three injections at two-week intervals.

**(i) D5901Fab:** Mouse monoclonal antibodies against OxlT were selected as previously described ^45^. Antibody-producing hybridoma cell lines were generated using a conventional fusion protocol. Hybridoma clones producing antibodies that recognised conformational epitopes in OxlT were selected by a liposome enzyme-linked immunosorbent assay (ELISA) on immobilised phospholipid vesicles containing purified OxlT, allowing positive selection of the antibodies that recognised the native conformation of OxlT. Additional screening for reduced antibody binding to SDS-denatured OxlT was used for negative selection against linear epitope-recognising antibodies. Stable complex formation between OxlT and each antibody clone was checked with fluorescence-detection size-exclusion chromatography.

Whole IgG molecules, collected from the large-scale culture supernatant of monoclonal hybridomas and purified using protein G affinity chromatography were digested with papain, and Fab fragments were isolated using HiLoad 16/600 Superdex200 gel filtration followed by protein A affinity chromatography. The sequence of the Fab was determined via standard 5’-RACE using total RNA isolated from hybridoma cells.

**(ii) 20D033Fv:** Single-chain Fv (scFv) fragments against OxlT were screened out from an immunised mouse phage displayed antibody library ^46^. Immunised mice were euthanised, and their splenocyte RNA isolated and converted into cDNA via reverse-transcription PCR. The V_L_ and V_H_ repertoire was assembled via an 18-amino acid flexible linker and cloned into the phage-display vector pComb3XSS. Biotinylated proteoliposomes were prepared by reconstituting OxlT with a mixture of egg PC and 1,2-dipalmitoyl-sn-glycero-3-phosphoethanolamine-N-(cap biotinyl) (16:0 biotinyl Cap-PE; Avanti), and used as binding targets for scFv-phage selection. Targets were immobilised onto streptavidin-coated paramagnetic beads (Dynabeads) or streptavidin-coated microplates (Nunc). After four rounds of biopanning, liposome ELISAs were performed on periplasmic extracts of individual colonies. Positive clones were collected and evaluated using a Biacore T100 (GE Healthcare).

Antibody scFv fragments are undesirable for use as crystallisation chaperones because they can intermolecularly form domain-swapped dimers, and the dimer-monomer equilibrium may increase structural heterogeneity. Therefore, we used Fv fragments for crystallisation trials. The Fv fragment were expressed in *Brevibacillus choshinensis* using the iRAT system ^47^. Culture supernatant was adjusted to 60% ammonium sulphate saturation, and the precipitate was pelleted, dissolved in TBS buffer (10 mM Tris-HCl, pH 7.5, 150 mM and NaCl) and dialysed overnight against the same buffer. Dialysed proteins were mixed with Ni-NTA resin equilibrated with buffer B (10 mM Tris-HCl, pH 7.5, 150 mM NaCl and 20 mM imidazole).

Bound proteins were eluted with buffer C (10 mM Tris-HCl, pH 7.5, 150 mM NaCl and 250 mM imidazole), mixed with TEV-His_6_ and dialysed overnight against TBS buffer. Cleaved His_6_ tag and TEV-His_6_ were removed using a HisTrap column equilibrated with buffer B. The tag-free Fv fragment was concentrated and loaded onto a HiLoad16/60 Superdex75 column (GE Healthcare) equilibrated with TBS buffer. Peak fractions were pooled, concentrated, flash frozen in liquid nitrogen and stored at −80 °C.

### Crystallisation

For crystallisation of oxalate-bound OxlT complexed with D5901-1A08-Fab, the purified OxlT was mixed with purified D5901-A08-Fab at a 1:1.3 molar ratio at 4 °C overnight and applied to a HiLoad 16/60 Superdex200 pg column (GE healthcare) using buffer D (20 mM MES-KOH, 200 mM potassium acetate, 10 mM potassium oxalate, 20% glycerol, and 0.51 mM DDM, pH 6.2) as running buffer. Purified sample was dialysed in buffer E (20 mM MES-KOH, 10 mM potassium oxalate and 0.51 mM DDM, pH 6.2). Crystals were obtained by the sitting-drop vapour diffusion method at 20 °C by mixing purified sample (∼10 mg/mL) with a reservoir solution of 0.1 M sodium citrate, pH 5.5, 0.05 M NaCl and 26% (v/v) PEG400. Crystals were frozen in liquid nitrogen in advance of data collection.

For crystallisation of ligand-free OxlT complexed with 20D033-Fv, purified OxlT was mixed with purified 20D033-Fv at a 1:2 molar ratio at 4 °C overnight and purified using Superdex200 Increase 10/300 GL (GE healthcare) in 20 mM MES-KOH, 10 mM potassium oxalate and 0.02% DDM, pH 6.2. Purified sample was reconstituted into a lipidic mesophase. The protein-LCP mixture contained 50% (w/w) protein solution, 45% (w/w) monoolein (Sigma) and 5% (w/w) cholesterol (Sigma). The resulting lipidic mesophase was dispensed as 50 μL drops into 96-well glass plates and overlaid with 0.8 μL of precipitant solution using an NT8-LCP crystallisation robot (Formulatrix) and were then covered with thin cover glasses.

Crystallisation setups and the 96-well glass sandwich plates (Molecular Dimension) were incubated at 20 °C. Crystals were obtained in a week under the following precipitation conditions: 100 mM Glycine, pH 9.0, 26–36% (v/v) PEG400, and 50–150 mM MnCl_2_. Crystals were harvested directly from the lipidic mesophase using Mesh Litholoops (Protein Wave) and flash cooled in liquid nitrogen.

### Data collection and structure determination

X-ray diffraction data for oxalate-bound OxlT and for ligand-free OxlT were collected at 1.0 Å at the SPring-8 beamline BL41XU using MX225HE (Raynoix) and BL32XU using an EIGER X 9M detector (Dectris, Ltd), respectively, under a cryostream operating at 100 K. Data were merged, integrated and scaled to 2.6 Å (oxalate-bound OxlT) and 3.1 Å (ligand-free OxlT) using the KAMO system ^48^, which exploits BLEND ^49^, XDS ^50^ and XSCALE ^51^ (Extended Data Table 1). Data were corrected for anisotropy using the STARANISO server ^52^. The correction deleted many weak reflections with very low spherical completeness in the higher resolution shells. For refinement, we used data to 3.0 Å (oxalate-bound OxlT) and 3.3 Å (ligand-free OxlT) that contained more than 25% (oxalate-bound OxlT) and 22% (ligand-free OxlT), respectively, of the data in the highest shell. The crystal structure was solved using molecular replacement with PHASER ^53^. The search models were structures of N- and C-terminal halves of the glycerol-3-phosphate transporter GlpT (PDB ID: 1PW4) ^19^ and an Fab fragment (PDB ID: 1XF4) ^54^ for oxalate-bound OxlT, and structures of N- and C-terminal halves of oxalate-bound OxlT determined in this study (residues 11–199 and 204–404, respectively) and a scFv fragment (PDB ID: 5B3N) ^55^ for ligand-free OxlT. Structure models were manually rebuilt with COOT ^56^ and refined with Phenix ^57^. In the ligand-free OxlT crystal, two units of OxlT (chain A and D) were found in an asymmetric unit. No significant structural difference was observed between the two (Cα RMSD of 0.365 Å for residues 15–410). Data collection and refinement statistics are shown in Extended Data Table 1. Ramanchandran statistics analysed with MolProbity ^58^ were 97.8% favoured, 2.2% allowed and 0.0% outliers for oxalate-bound OxlT, and 97.5% favoured, 2.5% allowed and 0.0% outliers for ligand-free OxlT.

### Transport assay

Transport activity of OxlT was evaluated by coupling with light-driven inward proton transfer by xenorhodopsin from *Rubricoccus marinus* (*Rm*XeR) co-expressed in *E. coli* ^37^. *E. coli* BL21 (DE3) cells were transformed with pRSF-OxlT ^37^ and pET21a-*Rm*XeR ^36^ and were cultured in LB medium containing 100 µg/mL carbenicillin and 50 µg/mL kanamycin at 37 °C. For the mutant OxlT assays, mutations were introduced into the RSF-OxlT vector via PCR using PrimeSTARMax (Takara Bio). Protein production was induced by adding 1 mM IPTG and 10 µM all-*trans* retinal (Sigma) at an absorbance of 0.8–0.9 at 600 nm. After culture at 20 °C for 20 h, *E. coli* cells were collected by centrifugation (3,500 × *g* for 5 min) and suspended with 50 mM K_2_SO_4_ to a cell density at 660 nm of ∼10.

The light-induced pH change of the cell suspension was monitored with a pH electrode (LAQUA F-72 pH metre, HORIBA) at 25 °C using continuous stirring. The cell suspension was first placed in the dark until the pH of the sample stabilised. The sample was then illuminated using a Xe lamp (MAX-303, Asahi Spectra) through a Sharp Cut Filter Y44 (a longpass filter at ≥ 420 nm, HOYA) for 10 min, and the pH change in the absence of oxalate (ΔpH_0_) was monitored. The light intensity was adjusted to ∼150 mW/cm^2^ at 550 nm using an optical power metre (#3664, Hioki) and an optical sensor (#9742, Hioki). The illuminated sample was placed back in the dark and when the pH stabilised, 5 mM potassium oxalate was added to the sample to enable transport via OxlT for 10 min. The sample was again illuminated under the same condition as above, and the pH change (ΔpH_S_) was monitored. The transport activity was evaluated by the difference in pH change (ΔΔpH) between ΔpH_S_ and ΔpH_0_; this was further corrected by subtracting the background differential pH change (ΔΔpH) measured with *E. coli* expressing *Rm*XeR alone. The activities for each mutant were normalised by the corrected ΔΔpH and the relative expression level, analysed by western-blotting using Penta·His Antibody (QIAGEN), of wild-type OxlT measured on the same day of experiment. We also performed an assay for the Y150A mutant; however, this mutant affected the expression level of *Rm*XeR due to unknown reasons, and we therefore excluded the Y150A result from this paper.

### Molecular dynamics simulation

The OxlT crystal structures were used as initial structures, with missing residues at the central loop modelled with MODELLER ^59^. Protonation states were analysed using PROPKA 3.1 ^60, 61^, with the default parameter. Based on the analysis, Lys355 exhibits a deviated p*K*_a_ value of 7.00 in the outward-facing structure. This deviation was not observed in the occluded structure (p*K*_a_ value of 8.61). Thus, both protonation states for Lys355 were considered in the outward-facing state. The OxlT protein was embedded in the membrane using the Membrane Builder plugin in CHARMM-GUI ^62, 63^. A phosphatidylethanolamine bilayer with a length of 120 Å for x and y dimension was used. The protein-membrane system was solvated with TIP3P water molecules and 150 mM KCl. We replaced Cl^-^ with oxalate ions with AmberTools17 ^64^. The final MD system contained 146015 and 143611 atoms for the occluded and outward-facing OxlT system, respectively. MD simulations were then performed using NAMD 2.12 ^65^. The Amber ff14SB and Lipid14 forcefields were employed to describe the protein and the membrane, respectively ^66, 67^. The oxalate ligand in solution was described with parameters determined by the electronic continuum correction with rescaling (ECCR), based on Ab Initio Molecular Dynamic simulation, developed by Kroutil et al. ^41, 68^. However, the oxalate ligand in the binding site of OxlT was described with parameters determined by the original RESP scheme, considering that the protein environment differs from that of water solution. The MD system was set up with a minimisation for 10,000 steps, heated from 0 to 10 K with a step of 0.1 ns per degree in NVT ensemble, then 10 to 310 K in NPT with a step of 0.2 ns per 30 degree, and 10 ns of equilibration with NPT ensemble simulation at 310 K. Then, production runs of 1.0 and 1.7 μs in NPT conditions were performed for the occluded and outward-facing OxlT (for each protonation state of Lys355) system, respectively. A temperature of 310 K was maintained with the Langevin thermostat, with the pressure set to 1 atmosphere using the Nosé-Hoover Langevin piston. Periodic boundary conditions were applied, and long-range electrostatic interactions were treated by the particle mesh Ewald method with a real space cut-off of 12 Å and a switch function at 10 Å.

To establish the simulation system with formate, a carboxylate moiety of the oxalate, toward to the Lys355, in the oxalate-bound occluded structure was replaced by a hydrogen atom to generate an initial structure of the formate-OxlT complex. GAFF force field parameters ^69^ were used for formate. The same equilibration and production protocols as described above were performed. The full relaxation of the OxlT protein at the end of the equilibration step guarantees a good adjustment of the binding site for a smaller ligand as well as a realistic conformation for production runs.

The water density during the simulation was calculated by a module from MDAnalysis ^70^ after the protein was centred and superimposed.

### QM/MM calculation

Several QM/MM models were employed with the oxalate-bound structure to assess the relevance of the binding site environment for the internal conformation of oxalate. First, the oxalate ligand was assigned to the QM part while the whole protein was assigned to the MM part. Second, the first shell of residues that interact directly with the ligand, (Gln34, Tyr35, Tyr124, Arg272 and Lys355) were added to the QM part. Third, the second shell of the binding site (Tyr150, Trp324, Tyr328 and Trp352) were added to the QM part to build a full binding site environment surrounding the oxalate ligand. All the QM/MM calculations were performed with ONIOM ^71^, implemented in Gaussian 16 ^72^. The density functional theory (DFT) method ^73, 74^ was used to treat the QM region at the B3LYP/6-31+G(d,p) level of theory ^75, 76^, including Grimme’s dispersion correction with Becke−Johnson damping (D3BJ)^77^. The MM region of the system was described by the same force field as that in the MD simulations. The electronic embedding scheme was used such that the MM region polarises the QM electronic density. An explicit link atom was added between the *α* and *β* carbons for each residue located in the QM region to handle the covalent boundary between the QM and MM parts. Minima of the potential energy surface were confirmed by having no imaginary frequencies. Additional pure DFT calculation of oxalate ligand with fixed side chains of the binding site residues were performed with the same QM level of theory as in the QM/MM calculations. As for QM/MM calculations, optimised structures were true energetical minima without imaginary frequencies.

## Acknowledgements

We thank Yayoi Nomura, Yoshiko Nakada-Nakura, Yumi Sato for technical assistance in the generation of antibodies, and Drs. Kazuya Hasegawa, Hideo Okumura, Yoshiaki Kawano, and Kunio Hirata, SPring-8, for X-ray diffraction data collection support. The synchrotron radiation experiments were performed at the BL41XU and BL32XU of SPring-8 and, with approvals of the Japan Synchrotron Radiation Research Institute (JASRI) (Proposal No. 2012B1096, 2015A1080, 2015B2080). Computations were partially performed using Research Center for Computational Science, Okazaki, Japan. This work was financially supported by JSPS KAKENHI Grant Numbers JP20H03195 (to A.Y.), JP18H02415 (to K.O.), JP26440086 (to T.H.), and Research Fund from Koyanagi Foundation (to A.Y.), Takeda Science Foundation (to Ta.S.), the Platform Project for Supporting Drug Discovery and Life Science Research [Basis for Supporting Innovative Drug Discovery and Life Science Research (BINDS)] from AMED under grant no. JP20am0101079 (to S.I.). The authors would like to thank Enago (www.enago.jp) for the English language review.

## Author contributions

T.H. and A.Y. conceived the study. Te.S., M.Y., Ta.S., K.H. and N.N. performed protein purification. N.N., H.I., T.H., and S.I. performed antibody preparation. Te.S., M.Y., A.Y, T.H., Ta.S., and K.H. performed crystallization and X-ray data collection. Ta.S., T.H., M.H., and A.Y. performed the structure analysis. M.H., K.K., A.Y., and Y.S. performed the transport assay. T.J.L. and K.O. performed molecular dynamics and QM/MM simulations. T.T. performed a preliminary molecular dynamics simulation. A.Y., K.O., Ta.S., N.N., T.J.L., and M.H. wrote the paper, together with input from all of the other authors.

---

**Extended Data Fig. 1.**
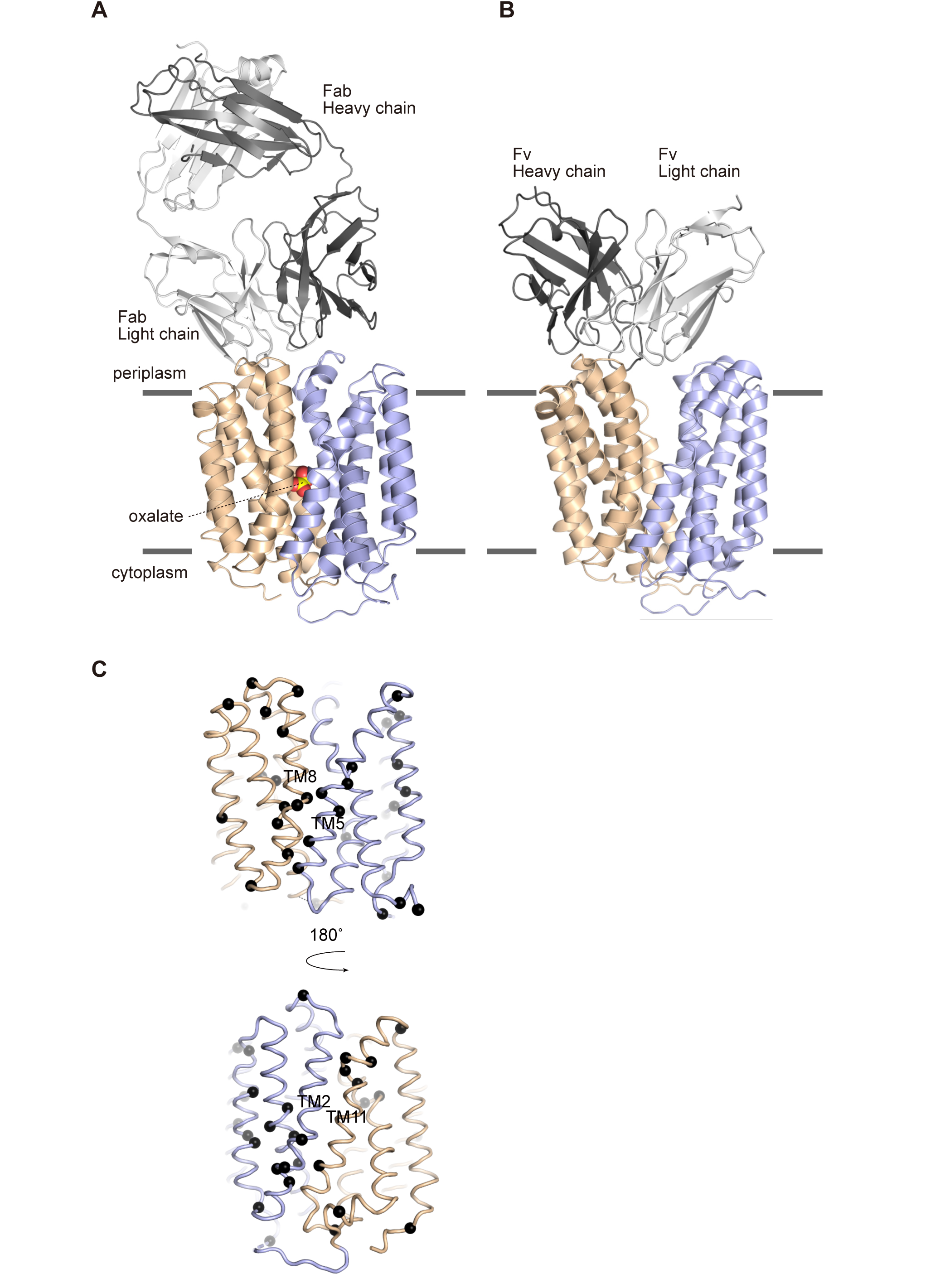
Structure of OxlT. (A) The oxalate-bound OxlT in complex with an Fab fragment (D5901Fab) and (B) the ligand-free OxlT in complex with an Fv fragment (20D033Fv). (C) Positions of the glycine residues mapped on the occluded OxlT structure, viewed from two different orientations. The black spheres indicate the Cα atoms of glycine residues.

**Extended Data Fig. 2.**
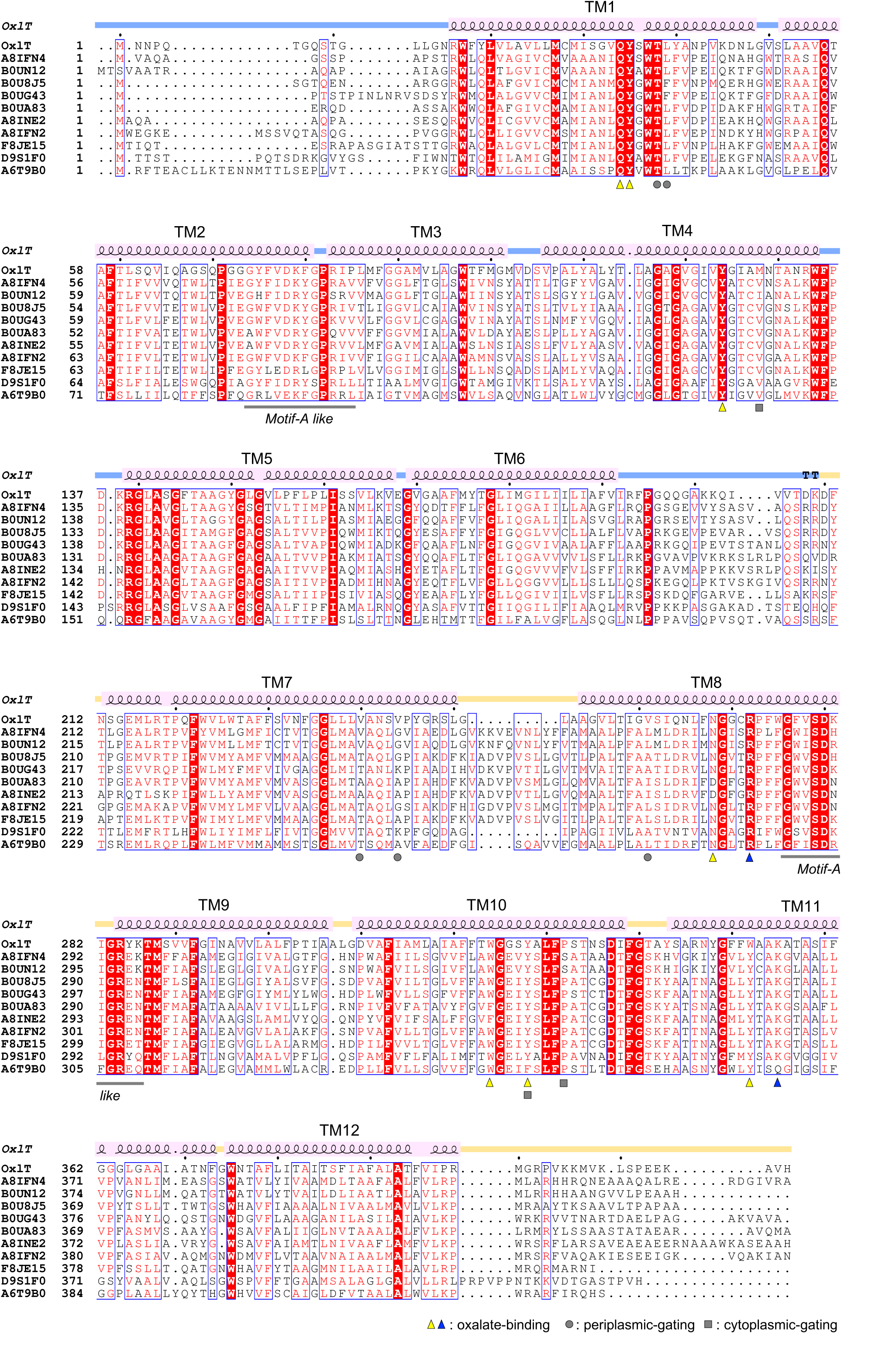
Amino acid sequence alignment of the OFA family proteins in MFS and the secondary structures of OxlT. Ten entries from oxalate/formate antiporter (InterPro ^43^ 026355) were aligned via the structure-based sequence using PROMALS3D ^78^; the alignment was drawn by the ESPript3.0 server (https://espript.ibcp.fr) ^79^. Note, all entries except of OxlT are uncharacterised.

**Extended Data Fig. 3.**
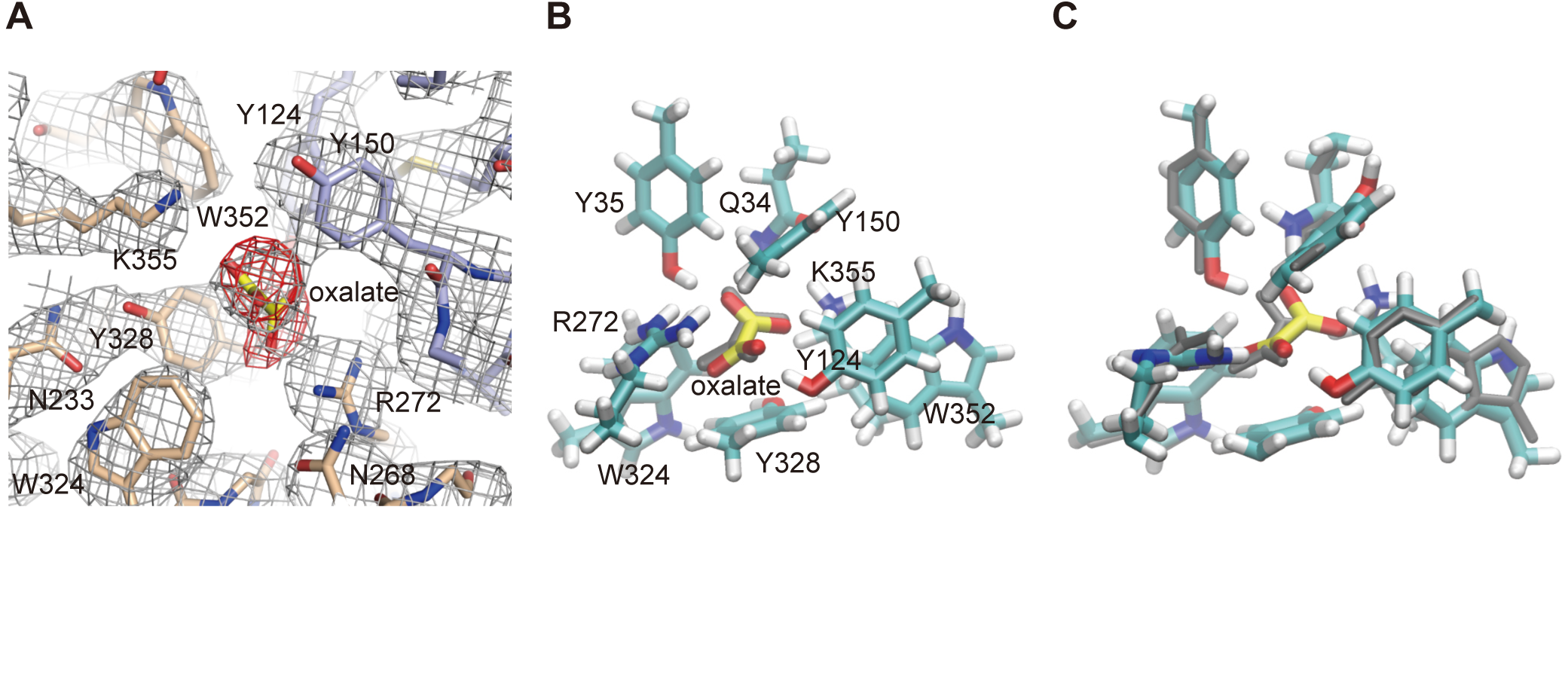
Oxalate binding at the occluded OxlT. (A) Simulated-annealing omit map of for the bound oxalate molecule, shown in red at 3.5 σ. 2*F*_o_-*F*_c_ map at the binding site in grey at 1.5 σ is also shown. (B) The QM-calculated structure of the oxalate binding site at the occluded OxlT. Binding site residues were set frozen, whereas the oxalate molecule was set free. The oxalate ion in the crystal structure is shown in grey. (C) The QM/MM-calculated structure of the oxalate binding site at the occluded OxlT. The QM/MM calculation was performed by applying the oxalate and neighbouring nine residues for QM and the other part of the transporter for MM calculation. Slight rearrangement of the residues in the binding site was observed from the crystal structure shown in grey, such as an additional H-bond formation with Gln34 and the oxalate.

**Extended Data Fig. 4.**
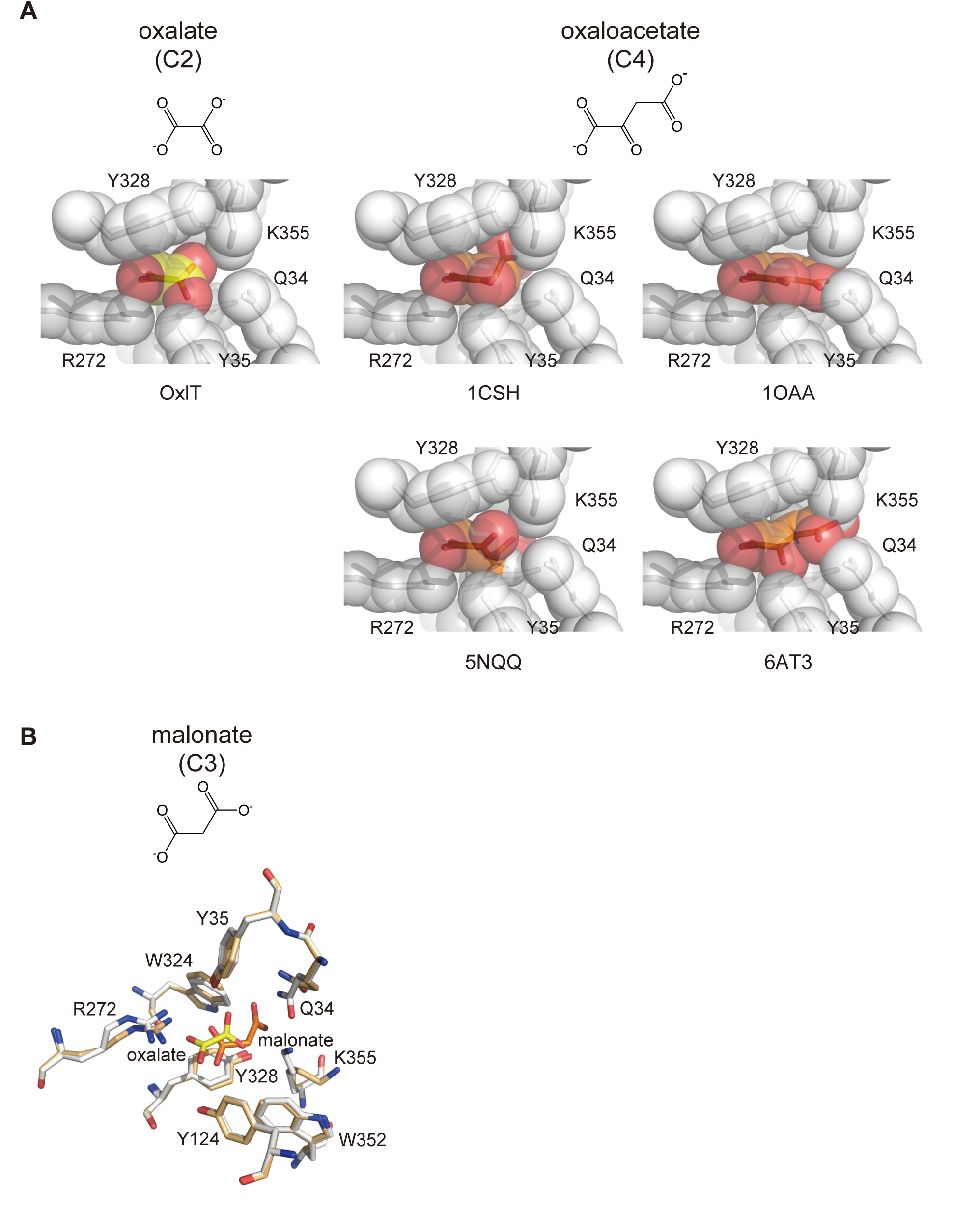
Presumed binding of different carboxylates at the substrate-binding site in OxlT. (A) The models of oxalate, the C4 dicarboxylate, and oxaloacetate, a C4 dicarboxylate intermediate in the Krebs cycle, bound to the occluded OxlT, coloured white. The oxaloacetate models with several different representative conformers observed in the PDB were simply supplanted with oxalate in the crystal structure; van der Waals surfaces are shown as spheres. (B) Docking model of the malonate-bound OxlT (OxlT, light orange; malonate, orange), superposed to the crystal structure of the oxalate-bound OxlT (OxlT, white; oxalate, yellow).

**Extended Data Fig. 5.**
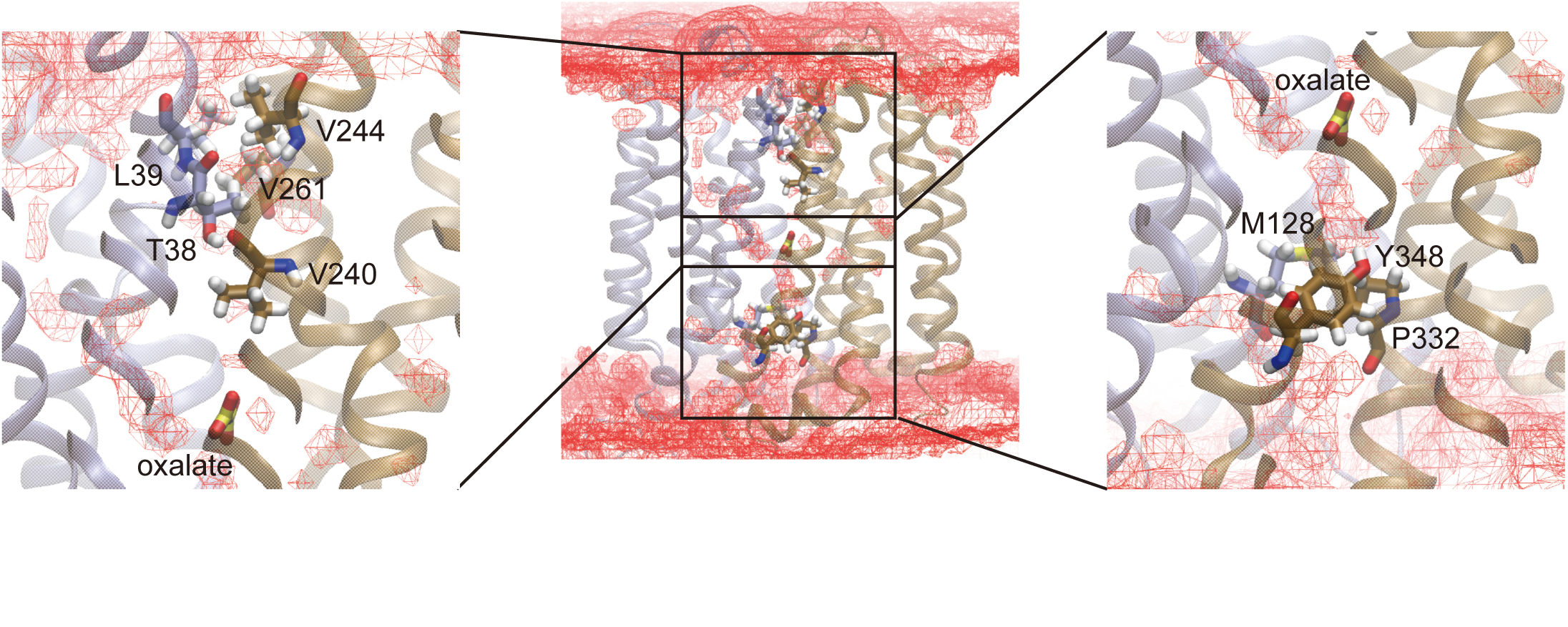
Water density in the occluded state during the simulation. The iso-surface of the relative density value of 0.5 to the bulk water is shown in red wires.

**Extended Data Fig. 6.**
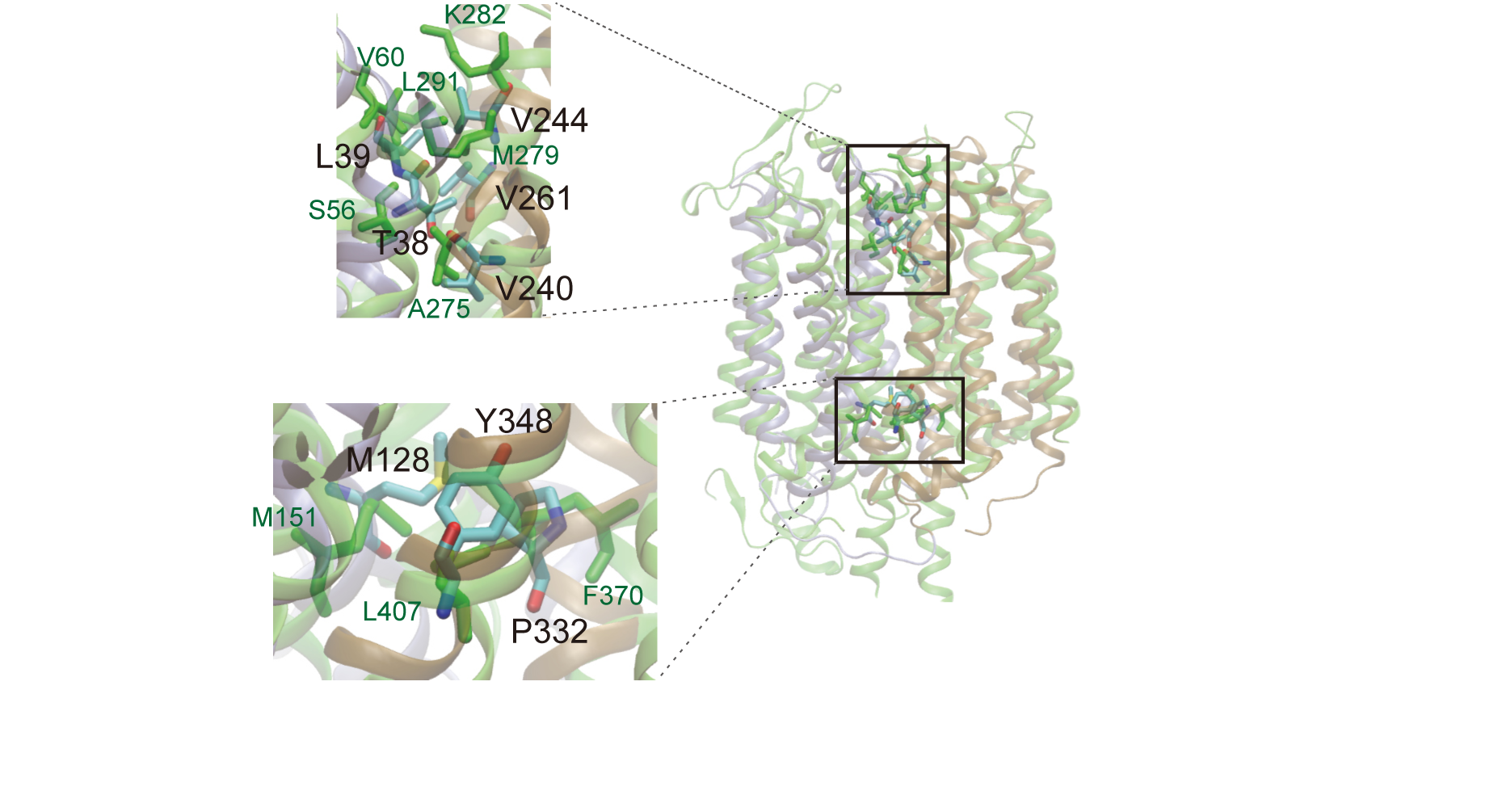
Structural alignment of OxlT and NarK. The NarK transporter is in green. Residues in OxlT and NarK located at similar positions are labelled in black and green, respectively. The NarK periplasmic gate consists of Ser56 and Val60 in TM1, Ala275 and Met279 in TM7 and Leu291 in TM8, whereas the NarK cytoplasmic gate consists of Met151 in TM4, Phe370 in TM10 and Leu407 in TM11 ^40^.

**Extended Data Fig. 7.**
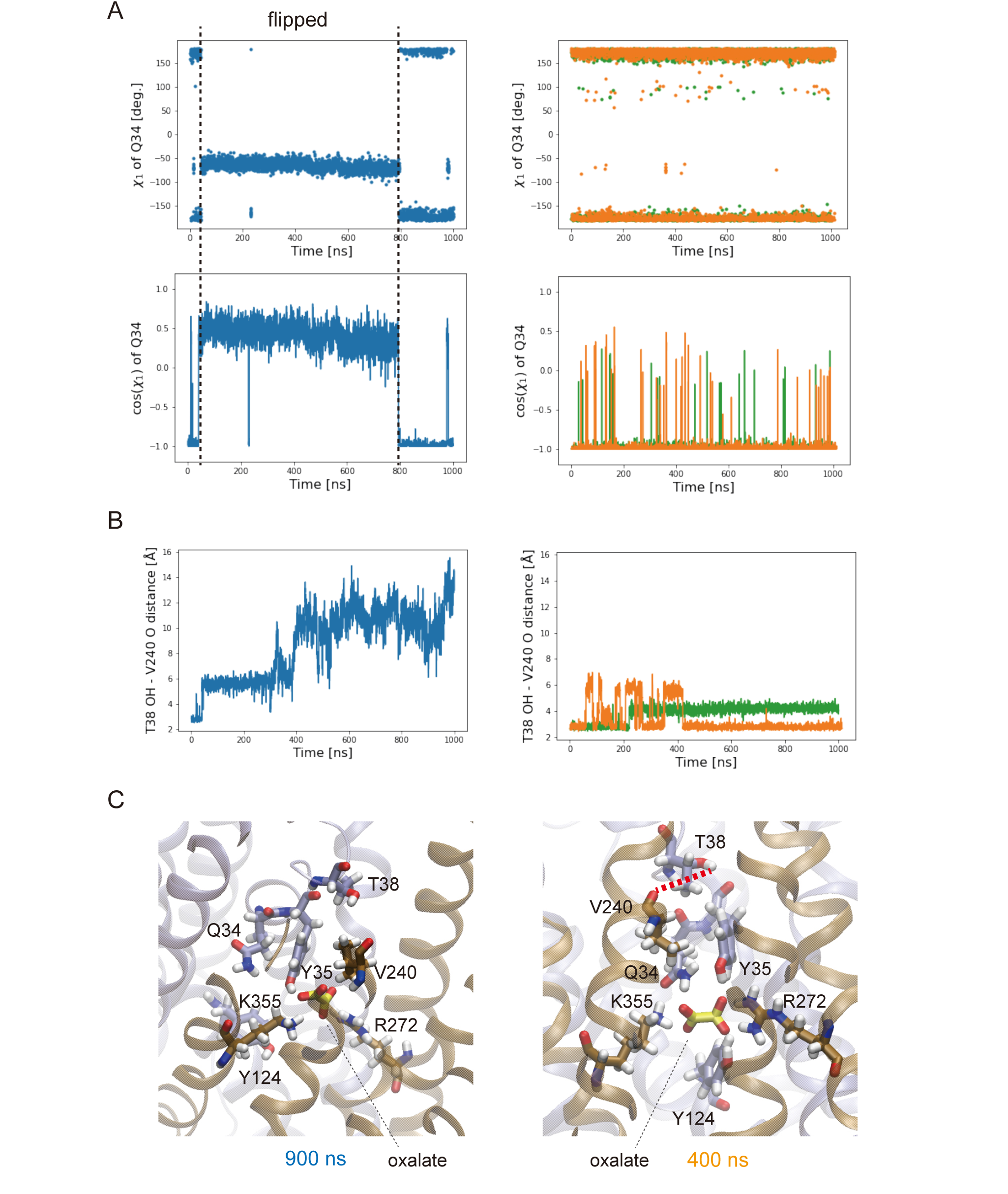
Gln34 side chain and hydrogen bond between Thr38 and Val240 in the simulation from the occluded conformation with oxalate. (A) The side chain dihedral x_1_ of Gln34 is shown for three independent trajectories with different colours. The flip of Gln34 side chain can be characterised by the change of the side chain dihedral χ. (B) The hydrogen bond donor and acceptor distance between Thr38 and Val240 is shown for three independent trajectories with different colours. (B) Snapshots of the binding site are shown. Data are derived from trajectories shown in Fig. 4D; the results from the trajectory with conformational transition (in blue) and the other two without conformational transition (in orange and green) are shown in the left and right subpanels, respectively.

**Extended Data Fig. 8.**
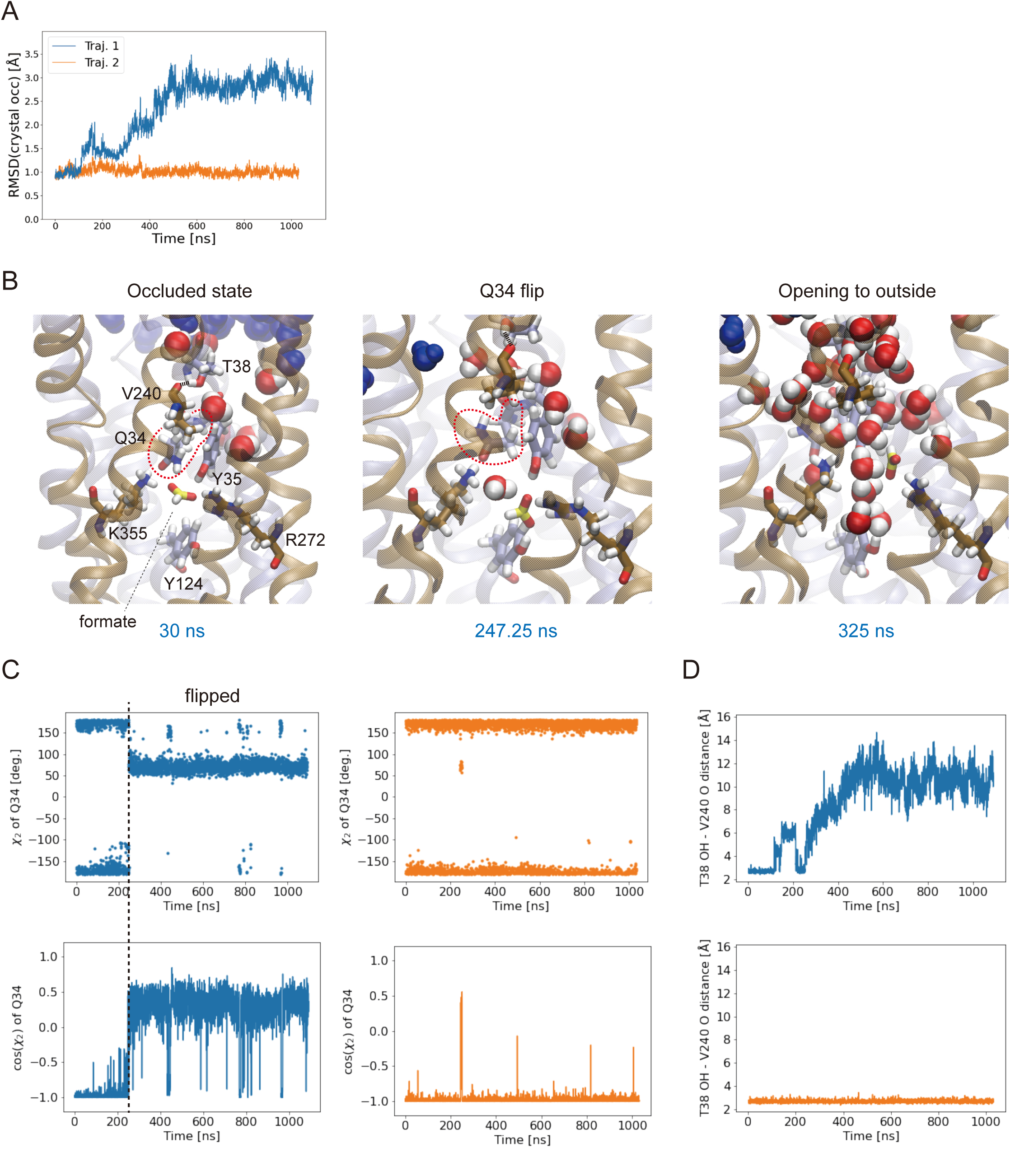
Simulation results from the occluded conformation with formate in the binding site. (A) RMSD plot for two independent trajectories. (B) Representative snapshots from a trajectory showing a transition from the occluded to the outward-open conformations. (C) The side chain dihedral χ_2_ of Gln34 is shown for two independent trajectories with different colours. Note that χ_1_ of Gln34 did not show a significant change upon the Gln34 flip in this case. (D) The hydrogen bond donor and acceptor distance between Thr38 and Val240 is shown for two independent trajectories with different colours.

**Extended Data Fig. 9.**
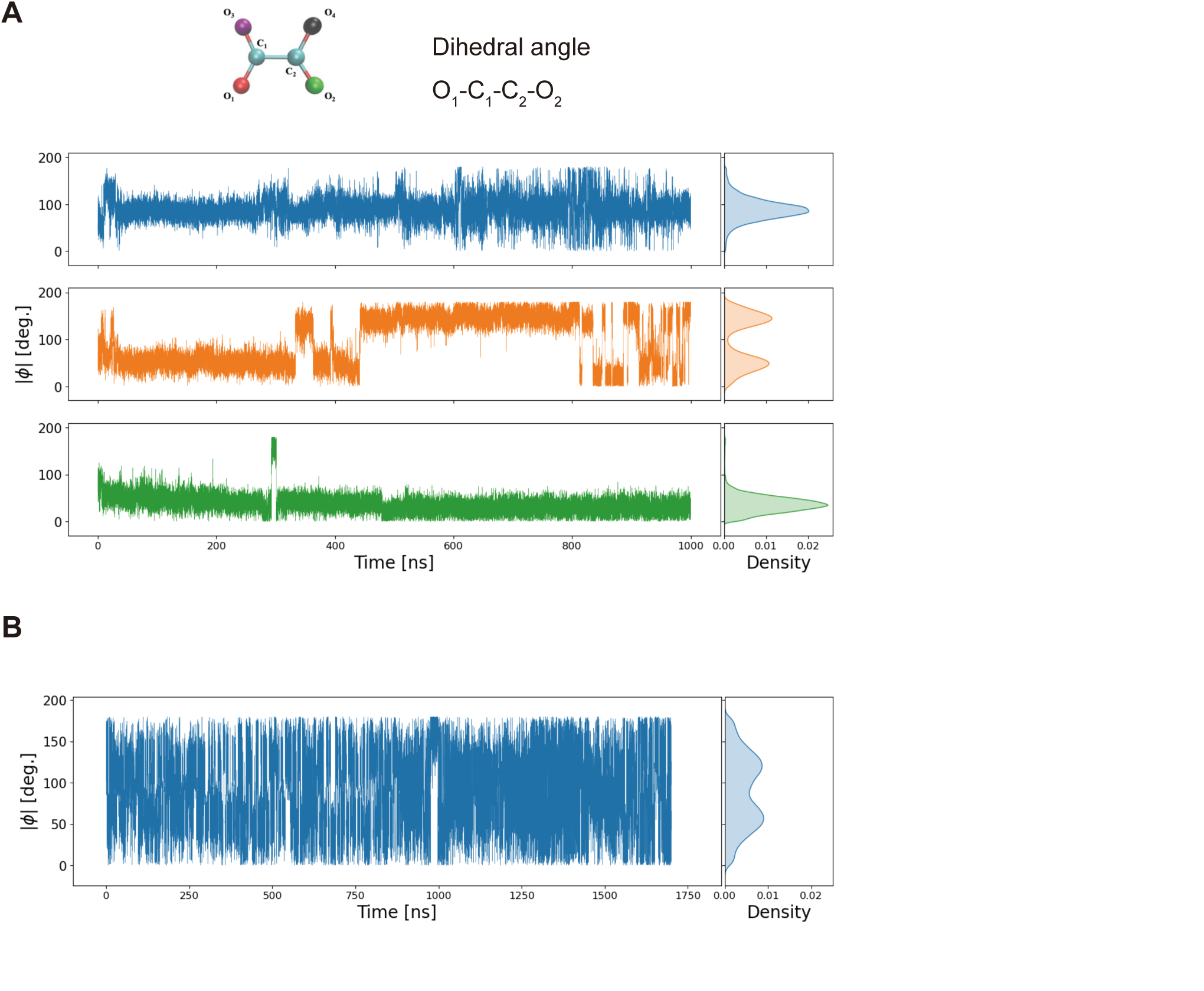
OxlT-bound oxalate conformation during MD simulations. (A) The dihedral angle of the oxalate ion in the binding site in MD simulations based on the oxalate-bound occluded OxlT crystal structure is shown for three independent trajectories in the same colour scheme as in Fig. 4D. (B) The dihedral angle of the spontaneously bound oxalate ion in the MD simulation started from the ligand-free outward-facing OxlT crystal structure with protonated Lys355 is shown.

**Extended Data Table 1.** X-ray crystallographic data collection and refinement statistics.

|  | **OxIT-Fab**<br>(PDB 7F7T) | **OxIT-Fv**<br>(PDB 7F7V) |
| --- | --- | --- |
| **Data collection** |  |  |
| Space group | $P2_12_12$ | $P2_1$ |
| Cell dimensions |  |  |
| *a*, *b*, *c* (Å) | 114.95, 233.19, 50.77 | 76.87, 181.67, 81.17 |
| $\alpha$, $\beta$, $\gamma$ (°) | 90, 90, 90 | 90, 111.37, 90 |
| Resolution (Å) | 46.64-2.60 (2.94-2.60)<sup>a</sup> | 47.26-3.10 (3.40-3.10) |
| *R*<sub>merge</sub> | 34.5(454.6)<sup>b</sup> | 83.9(332.3)<sup>b</sup> |
| *I*/$\sigma$(*I*) | 16.45 (1.71)<sup>b</sup> | 5.01 (1.26)<sup>b</sup> |
| *CC*<sub>1/2</sub> | 99.7 (65.1)<sup>b</sup> | 98.0 (68.6)<sup>b</sup> |
| Ellipsoidal Completeness | 92.9 (77.4)<sup>b</sup> | 88.4 (54.8)<sup>b</sup> |
| Spherical completeness | 52.4(8.8)<sup>b</sup> | 69.8(14.4)<sup>b</sup> |
| Redundancy | 99.8 (72.1)<sup>b</sup> | 24.8 (24.5)<sup>b</sup> |
| **Refinement** |  |  |
| Resolution (Å) | 21.08-3.00 (3.11-3.00) | 46.23-3.30 (3.42-3.30) |
| No. reflections | 20960 | 25465 |
| *R*<sub>work</sub> / *R*<sub>free</sub> | 23.7/27.8 (28.5/30.0) | 26.0/28.6 (38.9/39.2) |
| No. atoms |  |  |
| Protein | 6200 | 9517 |
| Oxalate | 6 | - |
| *B* factors |  |  |
| Protein | 51.2 | 53.9 |
| Oxalate | 42.4 | - |
| R.m.s. deviations |  |  |
| Bond lengths (Å) | 0.003 | 0.003 |
| Bond angles (°) | 0.59 | 0.61 |

<sup>a</sup>Values in parentheses are for highest-resolution shell.<sup>b</sup>Values reported by STARANISO anisotropy & Bayesian estimation server.

**Extended Data Table 2.** Oxalate dihedral angle and binding distances from the results of QM and QM-MM geometry optimisations.

|  | Dihedral angle | Binding distance |  |  |  |  |
| --- | --- | --- | --- | --- | --- | --- |
|  | O<sub>1</sub>-C<sub>1</sub>-C<sub>2</sub>-O<sub>2</sub><br>angle<br>(°) | O<sub>1</sub>-Y124(OH)<br>(Å) | O<sub>2</sub>-K355(NZ)<br>(Å) | O<sub>3</sub>-R272(NH2)<br>(Å) | O<sub>4</sub>-Y35(OH)<br>(Å) | O<sub>4</sub>-Q34(NE2)<br>(Å) |
| Crystal structure | 60.1 | 3.0 | 3.2 | 2.6 | 2.3 | > 3 |
| QM<br>9 frozen residues (Q34, Y35, Y124, Y150, R272, W324, Y328, W352, K355) + free oxalate |  |  |  |  |  |  |
| B3LYP | 68.2 | 2.7 | 2.8 | 2.8 | 2.6 | > 3 |
| B3LYP-D3BJ | 68.2 | 2.7 | 2.8 | 2.8 | 2.6 | > 3 |
| QM-MM<br>Oxalate, Q34, Y35, Y124, R272, K355 (QM) + other region of the protein (MM) |  |  |  |  |  |  |
| B3LYP-D3BJ | 50.2 | 2.7 | 2.6 | 2.6 | 2.8 | 2.8 |
| QM-MM<br>Oxalate, Q34, Y35, Y124, Y150, R272, W324, Y328, W352, K355 (QM) + other region of the protein (MM) |  |  |  |  |  |  |
| B3LYP-D3BJ | 52.3 | 2.7 | 2.7 | 2.6 | 2.7 | 2.8 |

